# Adaptive evolution in DNMT2 supports its role in the dipteran immune response

**DOI:** 10.1101/2020.09.15.297986

**Authors:** Tamanash Bhattacharya, Danny W. Rice, Richard W. Hardy, Irene L.G. Newton

**Affiliations:** Department of Biology, Indiana University Bloomington, USA

**Keywords:** Methyltransferase, Adaptive Evolution, Diptera, Drosophilidae, Culicidae, Virus, Wolbachia

## Abstract

Eukaryotic nucleic acid methyltransferase (MTase) proteins are essential mediators of epigenetic and epitranscriptomic regulation. DNMT2 belongs to a large, conserved family of DNA MTases found in many organisms, including holometabolous insects like fruit flies and mosquitoes, where it is the lone MTase. Interestingly, despite its nomenclature, DNMT2 is not a DNA MTase, but instead targets and methylates RNA species. A growing body of literature suggest DNMT2 mediates the host immune response against a wide range of pathogens, including RNA viruses. Evidence of adaptive evolution, in the form of positive selection, can often be found in genes that are engaged in conflict with pathogens like viruses. Here we identify and describe evidence of positive selection that has occurred at different times over the course of DNMT2 evolution within dipteran insects. We identify specific codons within each ortholog that are under positive selection, and find they are restricted to four distinct domains of the protein and likely influence substrate binding, target recognition, and adaptation of unique intermolecular interactions. Additionally, we describe the role of the Drosophila-specific host protein IPOD, in regulating the expression and/or function of fruit fly DNMT2. Finally, heterologous expression of these orthologs suggest that DNMT2’s role as an antiviral is host dependent, indicating a requirement for additional host-specific factors. Collectively, our findings highlight the adaptive evolution of DNMT2 in Dipteran insects, underscoring its role as an important, albeit non-canonical, regulator of host-pathogen interactions in mosquitoes and fruit flies.

## Introduction

Cellular DNA and RNA methyltransferases (MTases) are key mediators of epigenetic and epitranscriptomic regulation in eukaryotes. The former is carried out by a conserved family of DNA cytosine methyltransferases (DNMTs). The DNMT family includes true DNA MTases like DNMT1, DNMT3A, DNMT3B and DNMT3L (Goll and Bestor 2005; Denis, et al. 2011). The remaining member of the DNMT family is DNA MTase 2, or DNMT2, which, despite its name and sequence similarity to other DNMTs, has been demonstrated to have only residual DNA methylation activity *in vitro*. Instead, it has been shown that DNMT2 binds and methylates RNA substrates *in vivo* and *in vitro*, thus classifying it as a novel class of DNA-like RNA MTases (Jurkowski, et al. 2008; Denis, et al. 2011; Jeltsch, et al. 2017). Homologs of DNMT2 are present in the vast majority of animal, fungal and plant species. Notably, DNMT2 is the only known DNMT present in dipteran insects like *Drosophila melanogaster, Aedes aegypti, Aedes albopictus, Culex quinquefasciatus* and *Anopheles gambiae* (Lewis, et al. 2020). By extension, it is conceivable that all members of Drosophila and Culicidae families are DNMT2-only organisms.

Consistent with DNMT2’s role as a bona fide RNA MTase, evidence of genome-wide CpG methylation is nearly absent in these insects, leaving the biological role of this MTase unclear (Takayama, et al. 2014; Lewis, et al. 2020). Past studies investigating the biological function of DNMT2 suggest that it functions as a predominantly cytoplasmic protein during cellular stress, which can lead to increased longevity and greater host survival under stress conditions (Lin, et al. 2005; Schaefer, et al. 2010). Under these conditions DNMT2 is responsible for methylating transfer RNAs (e.g. tRNA_ASP,_ tRNA_GLU_), a modification that aids in protecting these RNA species from stress-induced degradation (Lin, et al. 2005; Schaefer, et al. 2010; Tuorto, et al. 2012). Aside from these known functions, the role of DNMT2 in the immune response is a fairly recent finding, following reports of its role in regulating the silencing of retrotransposons that otherwise contribute to cell stress (Phalke, et al. 2009; Schaefer and Lyko 2010; Durdevic, Hanna, et al. 2013; Durdevic and Schaefer 2013). Furthermore, proper functioning of DNMT2 in *Drosophila melanogaster* is required for efficient Dicer-2 activity and thus by extension, the RNA interference pathway (Durdevic, Mobin, et al. 2013). On its own, fruit fly DNMT2 inhibits several RNA viruses and protects the host against pathogenic bacteria (Durdevic, Hanna, et al. 2013; Bhattacharya, et al. 2017; Baradaran, et al. 2019). Furthermore, DNMT2 orthologs of several other arthropods have been shown to be involved in the colonization by pathogenic bacteria *(Helicoverpa armigera)*, RNA viruses *(Aedes aegypti, Aedes albopictus)* and Plasmodium *(Anopheles albimanus)*. Indeed, in previous studies we have demonstrated the roles of both *Drosophila melanogaster* and *Aedes* DNMT2 orthologs in regulating RNA virus infection. Notably, while DNMT2 in the fruit fly is responsible for limiting virus replication and production of infectious virus progeny, the *Aedes* orthologs seemingly play a proviral role in the mosquito host (Zhang, et al. 2013). Regardless, collectively these findings suggest that DNMT2 functions at the interface of host-pathogen interactions (Durdevic, Hanna, et al. 2013; Zhang, et al. 2013; Bhattacharya, et al. 2017; Baradaran, et al. 2019; Claudio-Piedras, et al. 2019).

Host genes involved in host immunity face strong selective pressure which is reflected in positive selection signatures in the genome e.g. Relish (Imd pathway) and Ci (Hedgehog signaling pathway) etc. (Sawyer, et al. 2003). This contributes to the adaptive evolution of these genes and the encoded products, driven by intermolecular interactions between the protein and its target e.g. pathogen associated molecular patterns (PAMPs). Given its recently identified role in arthropod immunity, we hypothesized that recurrent host-pathogen conflicts have impacted the molecular evolution of DNMT2 in Dipteran insects. In light of their well-documented history of harboring pathogens such as RNA viruses, we focused our analyses on members of Culicidae and Drosophila (Durdevic, Hanna, et al. 2013; Zhang, et al. 2013; Bhattacharya, et al. 2017; Claudio-Piedras, et al. 2019). Consistent with our hypotheses, we found significant evidence of positive selection along the ancestral lineage to all Dipteran DNMT2s as well as among DNMT2 orthologs of several members of the two aforementioned Dipteran families. Several amino acid positions in functionally important motifs of DNMT2 show evidence of positive selection. We found distinct differences in primary and tertiary protein structures between *Drosophila melanogaster* and *Aedes albopictus* DNMT2 that extend to other members of their respective families. We present evidence that regulation of DNMT2 is dramatically different in these two insects and that the antiviral function of DNMT2 is due to host cellular environment. Collectively, our results present evidence of adaptive evolution of DNMT2 in arthropods, underscoring its importance in host-pathogen interactions.

## Results

### Evidence of adaptive evolution in DNMT2

Prior studies have demonstrated that a high proportion of amino acid changes in Drosophila are driven by positive selection, and although statistical problems with models used to estimate positive selection may lead to “false positives” many studies, using different approaches, have detected a large proportion of positively selected sites in the Drosophila lineage, especially genes encoding for proteins that interact with pathogens (Sawyer, et al. 2003; Sella, et al. 2009; Jiang and Assis 2017; Kern and Hahn 2018). For example, Sawyer et.al found that a large majority (93%) of replacements present among 56 loci across *Drosophila melanogaster* and *Drosophila simulans* may be beneficial. As many dipterans are vectors for human pathogens, and infected by the endosymbiont *Wolbachia*, we hypothesized that DNMT2 in this group of insects may show evidence of positive selection. Indeed, evidence of positive selection in *Drosophila* DNMT2 has been reported earlier by Vieira et.al (Vieira, et al. 2018). However, that study was limited to identifying signatures of positive selection within *Drosophila* species. Here, our aim was to expand the scope of this previous analysis to additionally include DNMT2 orthologs from a total of 29 *Dipteran* insect species, which we evaluated for positive selection by maximum-likelihood analyses using CodeML (PAML package) (Yang 2007). Given the relevance of mosquitoes as disease vectors for viruses and other pathogens, our list included DNMT2 orthologs from a total of 20 species from the Culicidae family (Suborder: Nematocera), including 17 *Anopheles*, 2 *Aedes* and 1 *Culex* species (Figure 1A). Additionally, we included DNMT2 orthologs from 7 representative taxa spanning the Suborder Brachycera, including 5 members of the *Glossina* genus and one each from the following five genera: *Stomoxys, Musca, Drosophila* and *Phlebotomus*. DNMT2 orthologs from 6 non-dipteran insects were included as outgroups (Figure 1A). Consistent with our hypothesis, significant signatures of positive selection (raw p-value < 0.05) were detected along the branch ancestral to all Dipteran insects and *Phelebotomus papatasi* (Branches 2, 3). Additionally, we found significant signatures of positive selection along the ancestral branch leading to the entire Culicidae (Branch 19), as well as along relatively recent branches within that family and deeper branches (#19, 20, and 25; Figure 1A, Table 1). Notably, several branches to important mosquito taxa exhibited signatures of positive selection, including the *Culex quinquefasciatus* lineage (p=8e-6, Branch 54) *a*nd several *Anopheles* species or recently diverged internal branches: *Anopheles dirus, Anopheles minimus*, and branches 21, 30, and 42 (Figure 1A). Outside of the Culicidae family, signatures of positive selection were detected along lineages within the Brachycera Suborder of Dipteran insects (Figure 1A, Table 1). These included all ancestral lineages leading to genera within this Suborder, representing members of *Glossina* species, *Musca domestica, Stomoxys calcitrans, Phlebotomus*, and importantly, *Drosophila melanogaster* (Branches 2, 4, 5, 6, 7 and 10, Figure 1A). Additionally, in this analysis, the branch directly leading to *Drosophila melanogaster* was found to be under positive selection (Branch 5, Figure 1A). Taken together, these findings suggest an ongoing process of adaptive evolution in Dipteran DNMT2, suggesting potential roles of several, yet uncharacterized, DNMT2 orthologs in host-pathogen interactions. We have previously shown that Drosophila DNMT2 is antiviral and that *Wolbachia* infection modulates its expression (Bhattacharya et al., 2015). We therefore next aimed to perform more in-depth analyses of Drosophila DNMT2 orthologs to look for evidence of positive selection across 38 different Drosophila species encompassing the Sophophora (20 species) and Drosophila (18 species) sub-genera using CodeML (PAML package). DNMT2 sequence from *Scaptodrosophila lebanonensis* (Scaptodrosophila Genus) was used as an outgroup. The phylogenetic tree of Drosophila Dnmt2 orthologs inferred using Maximum-likelihood analyses was found to be largely congruent with previously reported phylogeny of Drosophila species, with distinct separation of DNMT2 orthologs into two known Drosophila subgroups (Figure 1B) (Russo, et al. 1995). Strong evidence of positive selection (raw p-value=0.002) was found in the lineage directly ancestral to all Sophophora (Branch 41) and weaker evidence (raw p-value=0.027) for the ancestral lineage to all Drosophila (Branch 2) and the lineages leading to *Drosophila grimshawi* (Branch 23), *Drosophila bipectinata* (Branch 64), *Drosophila fiscusphila* (Branch 66), and *Drosophila teissieri* (Branch 70). Notably, in this more focused analysis, positive selection was not found in *Drosophila melanogaster* (Branch 75), suggesting the absence of any recent adaptations since its divergence from other members of the Sophophora genus. Alternatively, we may lack statistical power to detect selection along these short branches. These findings suggest several instances of recent adaptive evolution within Drosophila DNMT2 since its divergence from Culicidae. Notably, these results are in line with the findings reported by Vieira et.al (Vieira, et al. 2018).

**Table 1.**
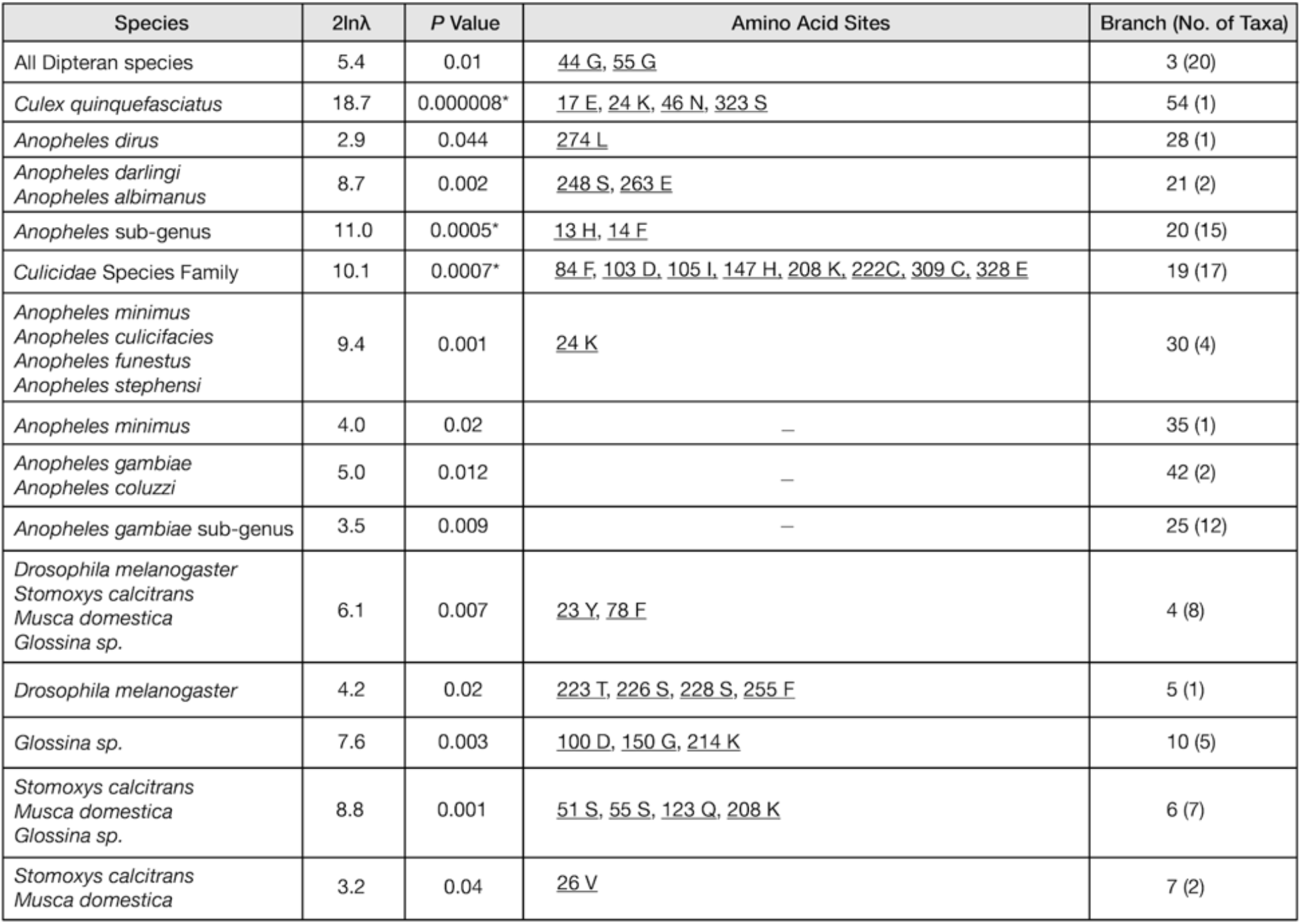
CodeML analyses result of positive selection among DNMT2 orthologs. Positively Selected Sites represent amino acid codon positions with ω > 1 (ω = d*N/*d*S*, BEB Posterior Probability > 0.80). Underlined codon sites represent those present in the ancestral lineage with BEB Posterior Probability > 95%.

**Figure 1.**
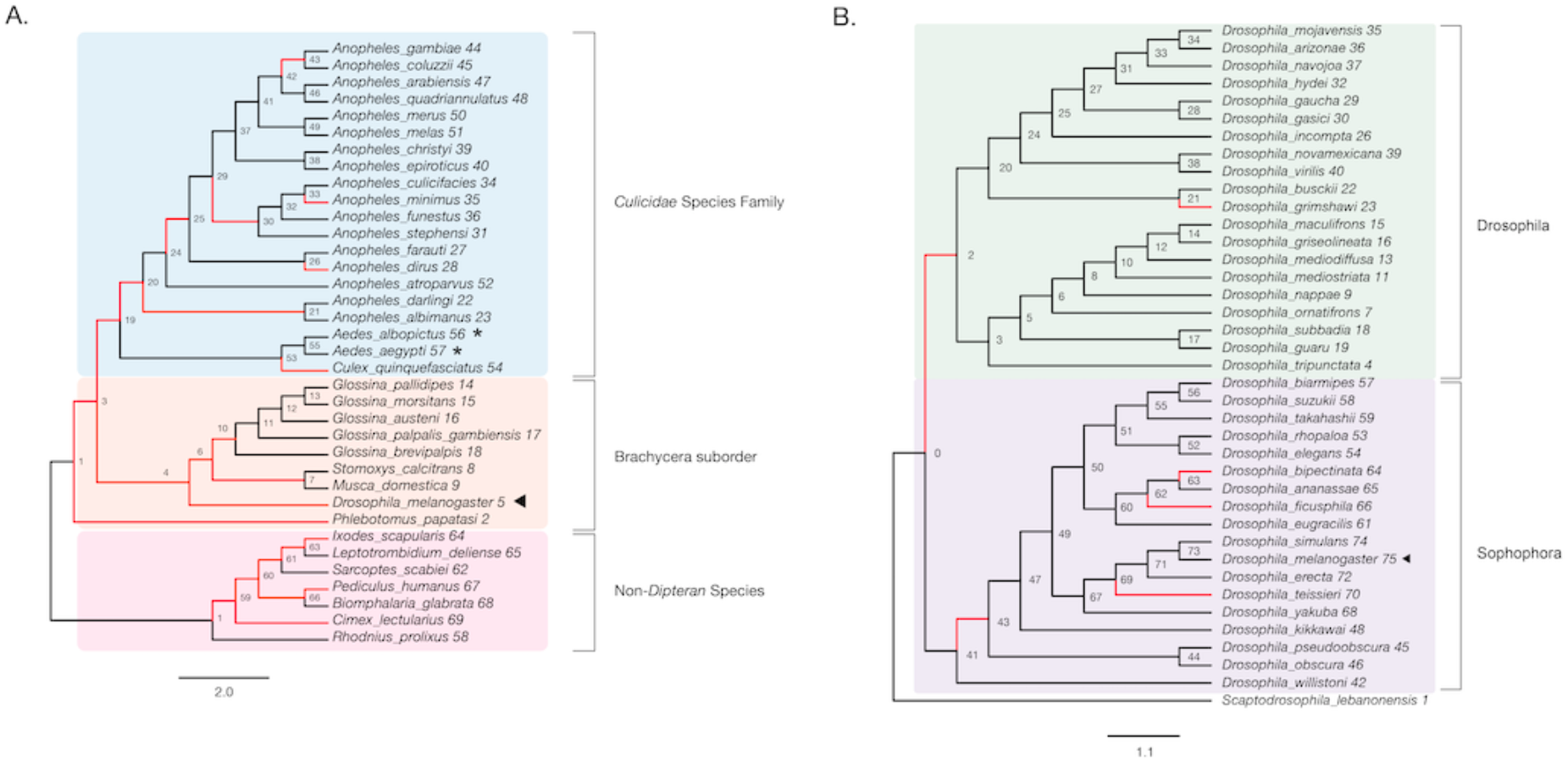
Evidence of adaptive evolution in DNMT2 orthologs. Branches numbered for reference in main text and Table 1. (A) Branch-site tests were conducted to detect positive selection (ω2 > 1, ω = d*N/*d*S*) across lineages of DNMT2 orthologs belonging to different species within the order Diptera (clades highlighted in light blue and orange for Culicidae family and the Brachycera suborder, respectively) and non-Dipteran (clades highlighted in light pink) animals. Maximum-likelihood (Deddouche, et al.) trees generated based on DNMT2 coding sequences using RAxML were used for the CodeML analyses. (A) Significant evidence (see Table 1 for details) of positive selection in DNMT2 is present along branches representing multiple insect species (ω2 > 1). These include several *Anopheles* and one *Culex* mosquito species, as well as several other Dipteran fly species Species whose DNMT2 ortholog(s) have been characterized as having anti-viral/microbial properties are indicated by black arrowheads, while those that have been described as having pro-viral properties are indicated by black asterisks. (B) Significant evidence of positive selection is present along the ancestral branch leading to the subgroup *Sophophora* (clades highlighted in light purple) and along branches leading to 4 *Drosophila* species. Taxa with adjacent black arrowheads represent DNMT2 ortholog with known anti-viral or anti-microbial activity. For both panels, branches under positive selection (ω2 > 1) are represented in red.

### Identification of codon sites under positive selection in DNMT2

The results from our previous CodeML analyses suggested multiple instances of positive selection along Dipteran lineages. To identify specific residues likely having undergone adaptive evolution, we used the Bayes Empirical Bayes (BEB) posterior probabilities from CodeML to identify amino acid sites having experienced positive selection (*d*_N_/*d*_S_ or ω>1) within the protein-coding regions of DNMT2. Notably, we found several sites from the ω>1 class with >95% probability across multiple Dipteran lineages (Table 1) and more specifically within the Drosophila genus (Table 2). Given their previous roles in host immunity, and the tractability of the model systems, we chose to focus our attention on codon sites present within lineages ancestral to or leading to *Aedes albopictus* DNMT2 (henceforth referred to as *Aa*DNMT2) and *Drosophila melanogaster* DNMT2 (henceforth referred to as *Dm*DNMT2.

**Table 2.**
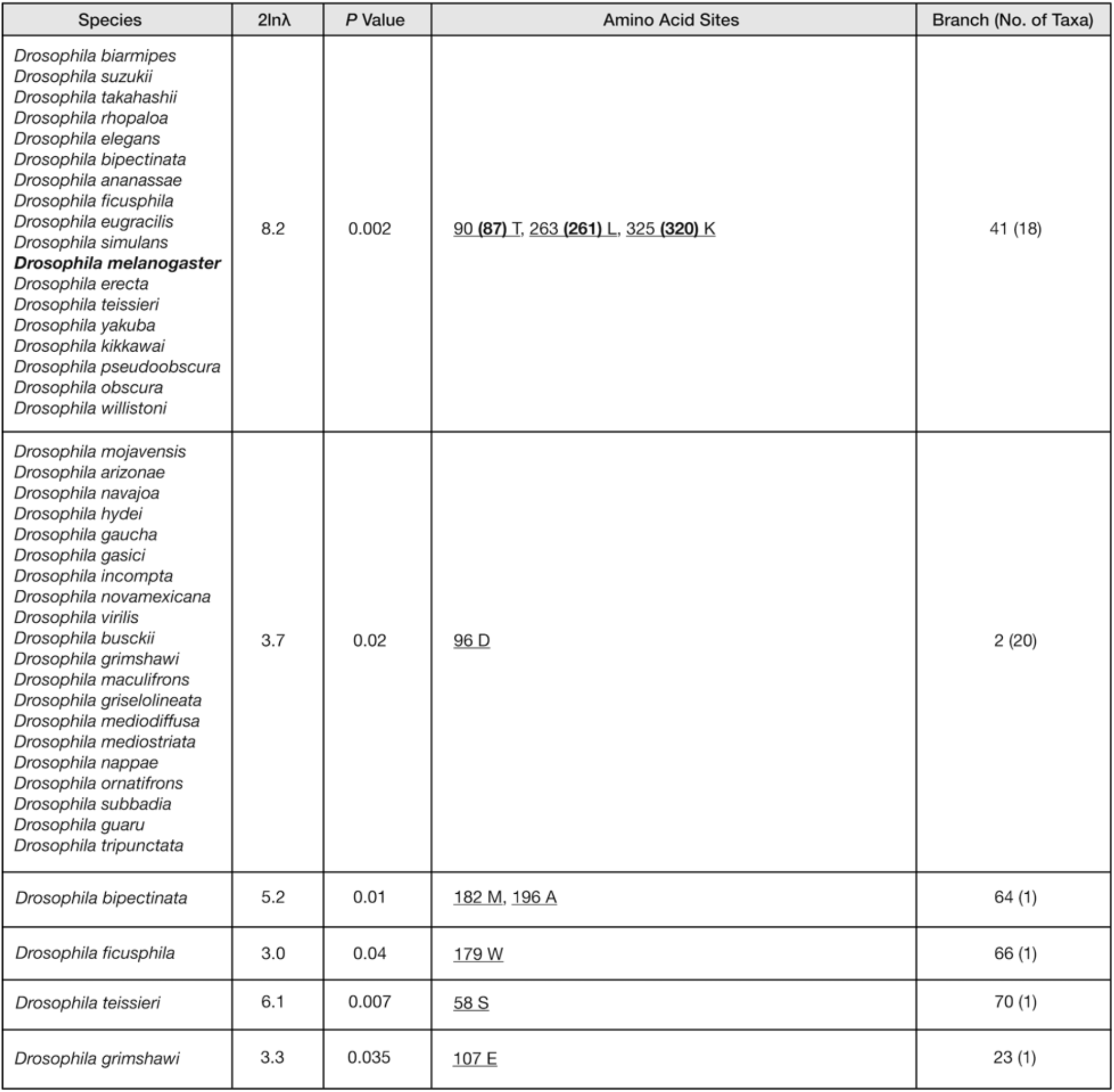
CodeML analyses result of positive selection among Drosophilid DNMT2 orthologs. Positively Selected Sites represent amino acid codon positions with ω > 1 (ω = d*N/*d*S*, <u>BEB Posterior Probability</u> > 0.95). *Drosophila melanogaster* taxa and associated amino acid sites are represented in bold. The codon sites within parenthesis relate to the positions for the same sites on the Dipteran multiple sequence alignment used in CodeML analyses for Table 1 and Figure 1A.

It is possible for changes identified along internal branches to have changed again later in some lineages. We looked at sites identified on internal branches to see which extant taxa still have them by assessing the degree of conservation at these sites within Culicidae and Drosophilidae families (Supplementary Figure 1). Two sites (44G, 55G), identified as being under selection among all Dipteran DNMT2s (Branch 3, Table 1) were found to be conserved in >80% of Culicidae and Drosophilidae species. Of the two sites, the ancestral variant 44G was conserved in the majority of the taxa (>83%, 24/30). In contrast, a conserved replacement site (55S) was found in the vast majority of species (>97%, 29/30), with only one *Anopheles* species harboring the ancestral 55G site. Aside from a few exceptions, conservation of the codon sites identified within Culicidae and Drosophilidae were limited to taxa within these respective families. Within Culicidae, our BEB analyses identified 19 amino acid positions under selection (Branches 54,19-21,28,30, Table 1). Mapping of these sites on a multiple sequence alignment of Culicidae species identified 4 amino acid sites unique to a single species, while the rest of the amino acid residues under selection were found to be present among multiple Culicidae taxa (Supplementary Figure 1A). Notably, despite the absence of selection detected along the *Aedes* lineage, 9 sites (Branch 19, Table 1) were found to occur within *Aedes* DNMT2 sequences, suggesting that these changes occurred prior to the divergence of this genus.

We next performed BEB analysis to identify codon sites under selection within *Drosophila* DNMT2. In order to represent all adaptive amino acid changes that have occurred in this taxa over its entire evolutionary period, 4 sites identified specifically in *Drosophila melanogaster* (Figure 1A, Branch 5, Table 1) were grouped alongside those identified in the most ancestral (2 sites) and most recent (2 sites) Dipteran lineages (Figure 1A Branches 3 and 4, Table 1), as well as sites identified in our *Drosophila* specific analyses (3 sites) appearing on the ancestral lineage to the Sophophora subgenus (Figure 1B, Table 2). Mapping of these 11 sites identified along lineages ancestral to *Drosophila melanogaster* revealed near-perfect conservation within Drosophila species from both Sophophora and Drosophila, suggesting that these changes occurred prior to the divergence of these subgroups. In contrast, sites identified along the branch ancestral to Sophophora were restricted to members of this subgroup (Supplementary Figure 1B). It should be noted that these 3 codon sites were identified previously by Vieira et.al, which adds support to our analyses (Table 2) (Vieira, et al. 2018). None of the 9 replacement amino acids unique to Drosophila were identified at the corresponding sites within members of the Culicidae, with one exception (Supplementary Figure 1B, Table 1-2).

We are ultimately interested in how these hypothesized adaptive changes in DNMT2 alter the function of the protein. Towards that end, we mapped these identified sites on the primary amino acid sequence of DNMT2 to determine their locations relative to previously identified functionally important regions (Falckenhayn, et al. 2016). Eukaryotic DNMT2 is broadly divided into two domains, the catalytic domain and the target recognition domain (TRD). The former can be further divided into ten functional motif regions (I – X) (Figure 2). Analyses of amino acid conservation across all sites between DNMT2 orthologs from Drosophila and Culicidae families suggest an overall 64% conservation in the primary amino acid sequence, with a higher degree of conservation, 77% within the catalytic region and 56% for the rest (Supplementary Figure 1). Of the 9 *Aedes* sites identified from the BEB analyses (Figure 1, branches 3 and 19), 5 were present within the catalytic domain (Supplementary Figure 1). These include one (55S) in the Motif II region, one (84F) in the active site loop adjacent to the catalytic PPCQ Motif IV region and two (323 S, 328 E) within the final Motif X region. One additional site (105I) was present within the catalytic domain albeit in a non-motif region. The rest of the 4 identified sites mapped to the TRD (Supplementary Figure 1A), suggesting that perhaps the *Aedes* DNMT2 has diverged in its target recognition. We next plotted the 11 sites from branches leading to D. melanogaster identified from our BEB analyses along the primary *Dm*DNMT2 amino acid sequence (Supplementary Figure 1B).

**Figure 2.**
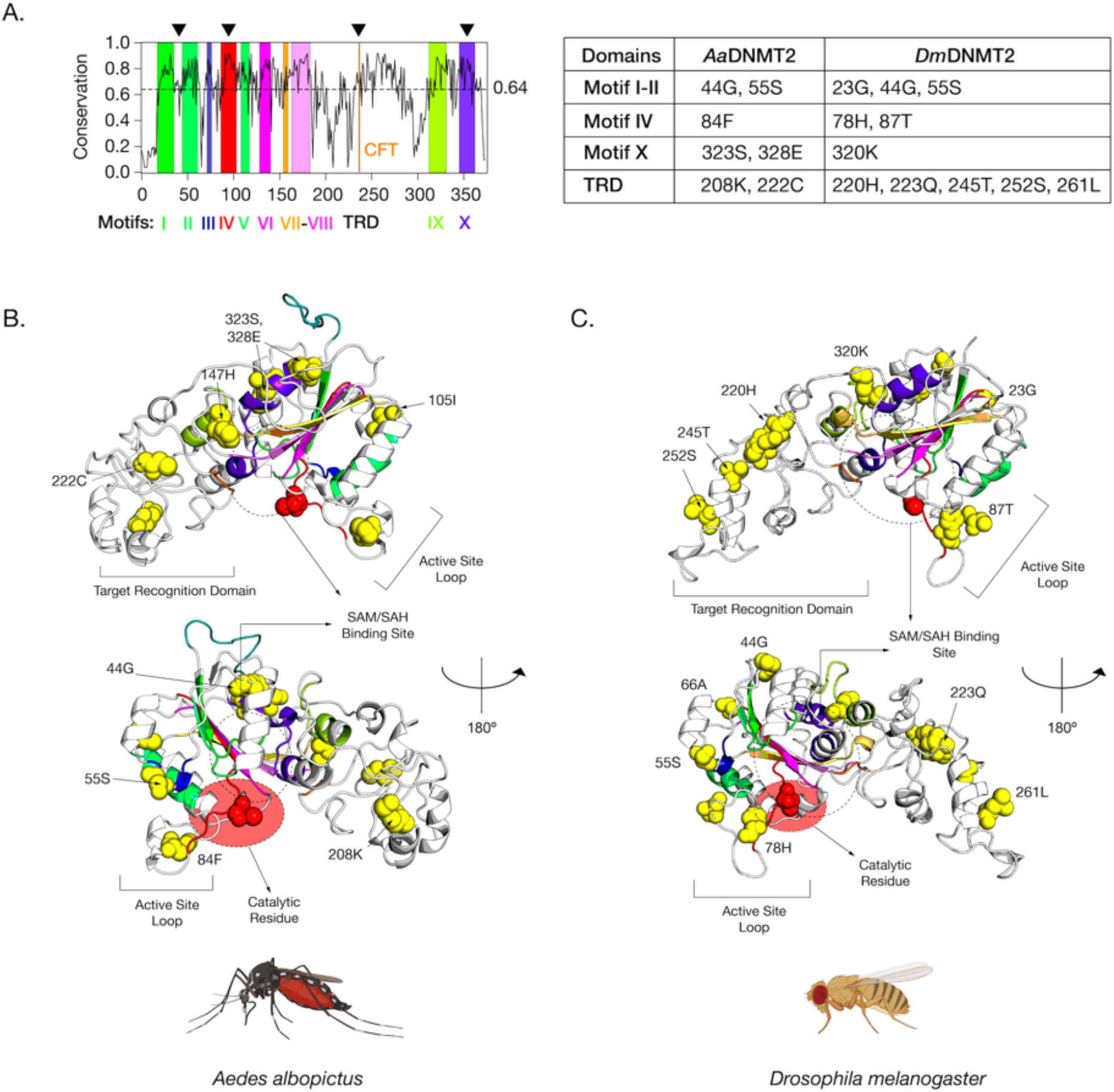
Amino-acid positions in *Drosophila melanogaster* DNMT2 potentially under positive selection. (A) Shannon conservation plot representing the degree of conservation (Y-axis) of DNMT2 orthologs present at every amino acid position (X-axis) across within DNMT2 orthologs from mosquitoes (Culicidae) and fruit flies (Drosophila). Colored boxes represent known DNMT2 functional motifs and domains involved in catalytic activity and target recognition (CFT). The mean conservation score (64%) across all amino acid positions is represented by the horizontal dotted line. Black arrows present on the top represent four major regions containing a majority of amino acid positions with evidence of positive selection and high posterior probability values (> 95%). These individual amino acids are also represented in the accompanying table to the right. (B, C) Spatial distribution of sites unique to each family are represented as yellow spheres on ribbon models of (B) *Aedes albopictus* (left, 9 sites) and (C) *Drosophila melanogaster* (right, 10 sites) DNMT2 structures visualized in PyMOL 2.4 (Schrödinger, LLC). The catalytically active cysteine residue (Cys, C) is represented in red. Predicted substrate i.e. S-adenosyl methionine (SAM) or S-adenosyl homocysteine (SAH) binding region is shown as a dashed oval. Functionally important active-site loop and target recognition domain are also indicated on each structure. The lower structures are rotated 180° relative to the upper ones.

While mapping the locations on the primary sequence allowed us to gauge the general location and conservation of these sites on the DNMT2 proteins of Culicidae and Drosophilidae, to assess the spatial importance of the amino acid sites identified in our BEB analyses with respect to MTase function, we next mapped a subset of the sites that were found within *Aedes albopictus* and *Drosophila melanogaster* on the 3D structures of *Aa*DNMT2 and *Dm*DNMT2, respectively (Figure 2). These orthologs were chosen as representative members of the Culicidae and Drosophilidae families, given their previously described roles in virus regulation and host immunity (Durdevic, Hanna, et al. 2013; Zhang, et al. 2013; Bhattacharya, et al. 2017; Claudio-Piedras, et al. 2019). Due to the absence of empirical structural information regarding *Aa*DNMT2 and *Dm*DNMT2, an intensive structural modeling approach using existing, experimentally solved crystal structures gathered from the Protein Data Bank (PDB) was used to generate predicted structures of these two DNMT2 orthologs (Figure 2). Furthermore, in order to gather a better understanding of the spatial distribution of the sites relative to the canonical MTase catalytic binding pocket, we used molecular docking to introduce the methylation substrate *S*-adenosyl-*L*-homocysteine (SAH) to identify the co-factor binding pocket (Grosdidier, et al. 2011).

Mapping of the aforementioned positively selected sites on *Aa*DNMT2 and *Dm*DNMT2 tertiary structures revealed the occurrence of positive selection at four major regions that were consistent between these two DNMT2 orthologs (Figure 2A). These include the four different regions: (1) region spanning Catalytic Motifs I and II (*Aa*DNMT2: 2 sites, 44G, 55S, *Dm*DNMT2: 3 sites, 23G, 44G, 55S), (2) Catalytic Motif IV Region and adjacent “active site loop” (*Aa*DNMT2: 1 site, 84F, *Dm*DNMT2: 2 sites, 78H, 87T), (3) Catalytic Motif X Region adjacent to the binding pocket for the canonical MTase co-factor *S*-adenosyl-methionine SAM and its resulting product *S*-adenosyl-homocysteine SAH (*Aa*DNMT2: 2 sites, 323S, 328E, *Dm*DNMT2: 1 site, 320K), (4) Target Recognition Domain involved in interactions with the nucleic acid target, facing away from the binding pocket, flanking the conserved CFT motif (*Aa*DNMT2: 2 sites, 208K, 222C, *Dm*DNMT2: 5 sites, 220H, 223Q, 245T, 252S, 261L) (Figure 2A). Incidentally, past studies indicate these four regions contribute significantly towards DNMT2’s MTase activity with regards to substrate binding and catalytic activity (Goll, et al. 2006). Furthermore, high clustering of sites in the TRD region is significant, given that they (*Aa*DNMT2: 208, *Dm*DNMT2: 261L) are located in a catalytically critical region that is known to penetrate the major groove of the nucleic acid substrate (Ye, et al. 2018). Finally, for both orthologs, a large proportion of sites present at the N-terminus (*Aa*DNMT2: 44G,55S,105I, *Dm*DNMT2: 23G,44G,55S,66A) and the TRD (*Aa*DNMT2: 147H, 222C, *Dm*DNMT2: 220H, 223Q, 245T, 252S) were found to be present on the solvent accessible surface (Figure 2B,C). These observations are in line with prior evidence that suggest that solvent exposure of protein surfaces have the strongest impact on adaptive mutations, likely driven by unique intermolecular interactions (Moutinho, et al. 2019). Indeed, we found this feature to not be limited just to *Aa*DNMT2 and *Dm*DNMT2, as mapping the positively selected sites on the tertiary structure of *Anopheles darlingi* DNMT2 revealed a vast majority of sites to occur on the solvent accessible protein surface (Figure 2B,C). Taken together, these observations suggest potential functional consequences of these amino acid substitutions on *Aedes albopictus* and *Drosophila melanogaster* DNMT2 with regards to catalytic activity and/or protein-protein interactions.

### *Aa*DNMT2 and *Dm*DNMT2 differ in structure

We and others have previously demonstrated regulation of RNA virus replication by *Aedes* DNMT2 orthologs in their respective host backgrounds, suggesting their involvement in host-pathogen interactions (Durdevic, Hanna, et al. 2013; Bhattacharya, et al. 2017). However, in contrast to the antiviral nature of *Dm*DNMT2, effects of *Aedes aegypti* (henceforth referred to as *Ae*DNMT2) and *Aedes albopictus* (*Aa*DNMT2) are distinctly proviral (Zhang, et al. 2013). Distinct molecular evolution between *Aedes* and *Dm*DNMT2 orthologs led us to next investigate whether potential differences in structure and/or regulation might contribute to the functional differences of these DNMT2 orthologs, using *Aa*DNMT2 and *Dm*DNMT2 as representative MTase orthologs from Culicidae and Drosophilidae families.

First, we assessed the broader differences in protein sequence across members of Culicidae and Drosophilidae. Multiple sequence alignment of DNMT2 primary amino acid sequences indicate that differences between overall fly and mosquito DNMT2 orthologs are most notable in the N-terminal end and the C-terminal (residues 282-292) target recognition domains. This is evidenced by the low (≤ 20%) conservation scores in these two regions (Figure 2A). The N-terminal end of mosquito DNMT2 is variable in length across different taxa in the Culicidae family and are, on average, 7-12 aa longer than the Drosophilidae counterparts, with the *Anopheles darlingi* DNMT2 ortholog being 47 aa longer in length (Supplementary Figure 3). In contrast to mosquito DNMT2, we found two instances of extended N-termini within Drosophilidae DNMT2; *Drosophila busckii* (17 aa) and *Drosophila serrata* (4 aa). We found an overall lack of sequence conservation among the different Culicidae orthologs, aside from a few residues that are conserved within members of the *Aedes* and *Anopheles* genus. *In silico* prediction analyses also showed this region to be devoid of any ordered secondary structure, suggesting conformational flexibility and potential to participate in protein-protein interactions. The other prominent difference in primary sequence between DNMT2 orthologs from these two Dipteran families occur within the target recognition domain (TRD), which is extended (10-12 aa) in the vast majority of Drosophilidae DNMT2 orthologs, with the exception of *Drosophila ananassae* and *Drosophila bipectinata* (Supplementary Figure 3). However, unlike the N-terminal extension within Culicidae, the extended TRD of Drosophilidae DNMT2 contains a conserved stretch of three residues (KSE) that constitute the start of a predicted α-helix (Figure 3, Supplementary Figure 3). Taken together, it is conceivable that such differences in the TRD contribute to differential substrate-MTase interactions between Culicidae and Drosophilidae DNMT2 orthologs.

**Figure 3.**
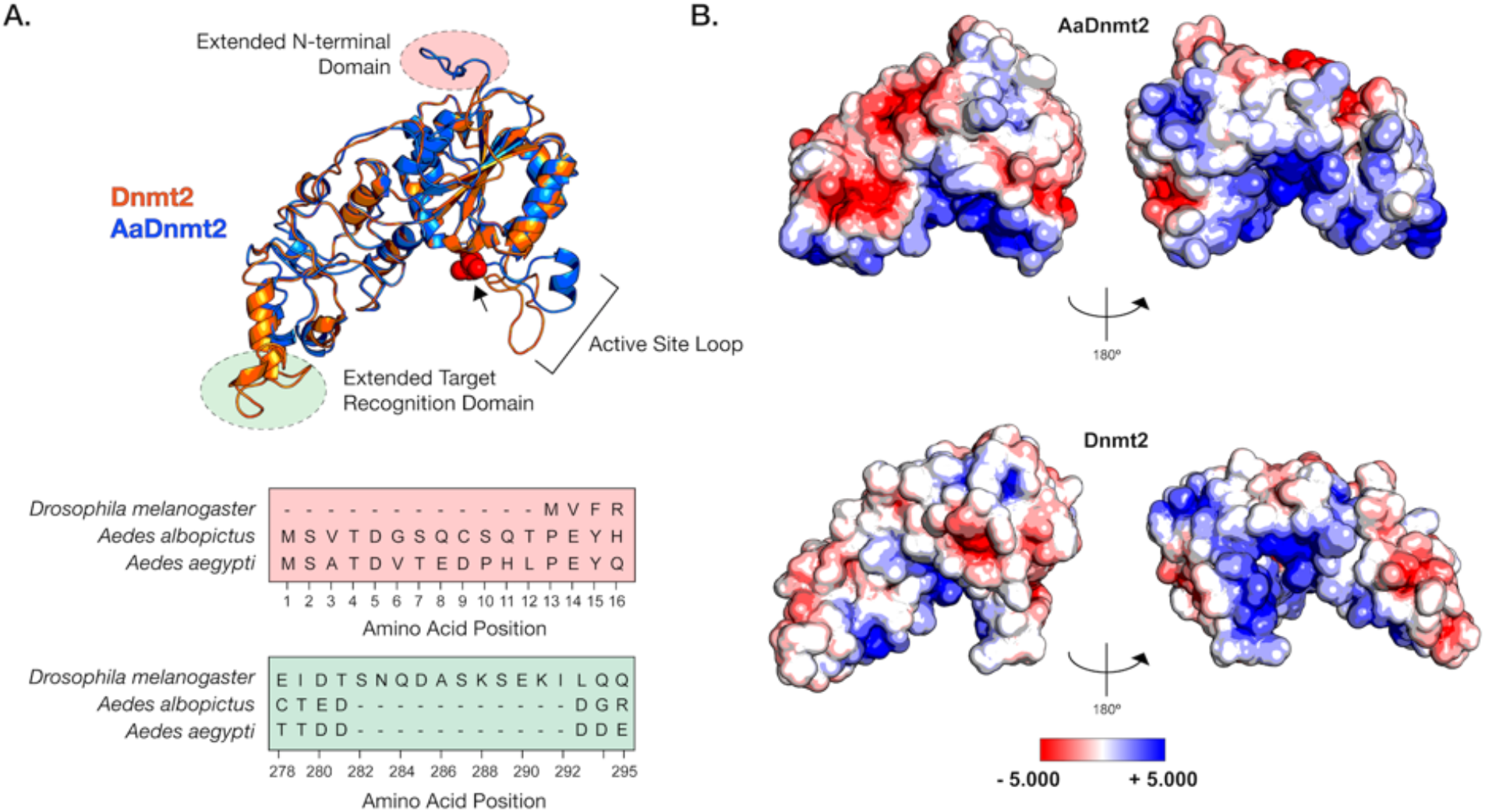
Structural differences between Drosophila and Aedes DNMT2 orthologs. Structures of DNMT2 orthologs from *Drosophila melanogaster* (*Dm*DNMT2) and *Aedes albopictus* (AaDNMT2) were generated using homology modelling (Phyre 2). (A) Superimposed ribbon diagrams of DNMT2 orthologs from *Drosophila melanogaster* (DNMT2, in blue) and *Aedes albopictus* (AaDNMT2, in orange) outline key structural differences. Primary sequence alignment of the two orthologs (46% overall amino-acid sequence identity) indicate significant differences in the N-terminal end (indicated in pale red on the ribbon diagram and the sequence alignment below) and the Target Recognition Domain (TRD) (indicated in pale green on the ribbon diagram. The catalytic cysteine residue (Cys 78) present within the highly conserved PP<u>C</u>Q motif is represented as red spheres). (B) Electrostatic Potential Surface Visualization models of DNMT2 orthologs were generated through PyMOL 2.4 (Schrödinger, LLC.) using the in-built Adaptive Poisson-Boltzmann Solver (APBS) plug-in. Colored scale bars indicate the range of electrostatic potentials calculated based on amino-acid compositions of each DNMT2 ortholog. The rotation symbol reflects structural features viewed 180° apart along the vertical axis.

*Aa*DNMT2 (344 aa) and *Dm*DNMT2 (345 aa) are comparable in size, sharing 46% amino acid sequence identity. However, while this does not necessarily imply that these DNMT2 orthologs differ to the same extent when it comes to their overall structure, in line with other Culicidae and Drosophilidae species, these orthologs exhibit major differences in two regions; the N-terminus and the Target Recognition Domain (Figure 2A). We therefore compared tertiary structures of these orthologs to identify how these differences affect their respective structures. The extended N-terminal end of *Aa*DNMT2 remained surface exposed in an unstructured, flexible conformation, indicating the ability to interact with potential interaction partners (Figure 3A). The extended TRD region within *Dm*DNMT2 was also found to be mostly surface exposed, adopting a short α-helical conformation at the C-terminal end. Comparison to crystal structures of DNMT2 from army worm *(Spodoptera frugiperda, PDB ID: 4HON)* and fission yeast (*Schizosaccharomyces pombe, PDB ID: 6FDF)* indicate that the rest of the TRD is unstructured and conformationally flexible. Given the importance of the conformation state of this TRD region for interactions with the nucleic acid substrate, the extended region within *Dm*DNMT2 carries the potential to alter MTase-substrate interactions (Goll, et al. 2006).

Outside of the two aforementioned regions, other notable structural differences are in the 20aa long active site loop region adjacent to the catalytic PPCQ motif. This region appears to be more structured in *Aa*DNMT2 relative to *Dm*DNMT2, consisting of a short stretch of residues forming an α-helix (Figure 3A). We found this feature to be consistent with the *in silico* secondary structure prediction for this *Aa*DNMT2 region. However, in contrast to the estimated 3D structure, this α-helical stretch was predicted to be extended for *Dm*DNMT2, spanning the entirety of the active site loop. This is likely a result of differences in the amino acid composition within this region between the two orthologs, where residues present within *Aa*DNMT2, e.g. Proline (P), Valine (V), Phenylalanine (F), are more likely to disrupt formation of α-helices. It should be noted, however, that this region has been suggested to adopt different structural conformations, switching between structured and unstructured α-helices, upon nucleic acid binding [34]. Multiple sequence alignment and structural modelling of Culicidae and Drosophilidae DNMT2 orthologs suggests that this feature is consistent within members of the respective families. At the same time, it should be noted that that these modelled structures are built on snapshots of otherwise dynamic crystal structures, and therefore limit our interpretation given that each structure is restrained to a singular, static conformation.

Aside from differences in secondary and tertiary structure, physiochemical properties of amino acids contribute to their spatial distribution and the propensity to remain either buried or exposed in a solvent accessible conformation. This attribute of proteins can also influence interactions with other biomolecules, which for enzymes like MTases include cognate interaction partners such as regulators and/or nucleic acid substrates. We therefore asked whether *Aa*DNMT2 and *Dm*DNMT2 differ significantly in terms of their surface charge distribution profiles. Mapping of electrostatic charge densities on solvent accessible 3D surfaces revealed an overall greater distribution of charged residues on the surface of *Aa*DNMT2. This included a distinctly larger patch of negatively charged residues in the TRD (Figure 3B). As expected, both DNMT2 orthologs contained a high density of positive charge in and around the catalytic region known to bind the negatively charged nucleic acid substrate. Additionally, in line with its role in substrate binding, the catalytic helix adjacent region of *Aa*DNMT2 was determined to be largely positively charged. This attribute however was noticeably absent from *Dm*DNMT2, whose extended catalytic helix adjacent region was found to be moderately negatively charged (Figure 3B).

Taken together, structural superposition of *Aa*DNMT2 and *Dm*DNMT2 demonstrates overall structural congruency between the two orthologs, but also shows significant differences which potentially indicate unique protein-protein and/or protein-substrate interactions for each ortholog.

### Drosophila IPOD regulates DNMT2 expression

Pathways and host factors involved in regulating DNMT2 expression in dipteran insects are poorly understood. In a past study, Kunert et.al. identified a potential host factor in *Drosophila melanogaster*, the aptly named Interaction Partner of DNMT2 or IPOD, in regulating *Dm*DNMT2 expression and function (Kunert 2005). However, it is unclear whether IPOD is involved in DNMT2 regulation within all Dipteran insects or whether distinct modes of DNMT2 regulation have evolved across different Dipteran families. In light of our previous results highlighting differences between Drosophilidae and Culicidae DNMT2, we next investigated the presence and conservation of IPOD orthologs among the species included in this study. Additionally, we examined the role of this protein in DNMT2 regulation within *Drosophila melanogaster*.

The protein IPOD is predominantly restricted to *Drosophila* species, based on BLAST searches of Dipteran insect genomes found to encode DNMT2 orthologs (Figure 4A, Supplementary Figure 4). Importantly, this was not due to the absence of available sequence information, as nearly complete genome assemblies are present for all taxa except for *Polypedilum vanderplanki*. Phylogenetic analyses of these *Drosophila* IPOD sequences illustrates its conservation within both Drosophila and Sophophora sub-groups of the Drosophila genus, with an average 46% amino acid sequence identity across all positions (Figure 4A,B). Furthermore, MirrorTree analyses of Drosophila DNMT2 and IPOD ortholog phylogenies revealed significant mirroring of the two trees, indicating a strong inter-protein co-evolutionary relationship between the two; Correlation: 0.787, P Value ≤ 0.000001 (Figure 4). This was further validated by the results from TreeCmp analyses assessing the Robinson Foulds (RF) and Matching Split (Williams, et al.) distances between Drosophila IPOD and DNMT2 trees, which showed a similar high congruence between the two tree topologies; RF (0.5) = 8, MS = 27.0. In contrast, very low congruence, with normalized distances ≤ 0.4 was found when the trees were compared to random trees generated according to Yule (RF/MS to YuleAvg) and uniform (RF/MS to UnifAvg) models; RF (0.5)_to UnifAvg = 0.3841, RF (0.5) to YuleAvg = 0.3852, MS_to UnifAvg = 0.2426, MS to YuleAvg = 0.2880.

**Figure 4.**
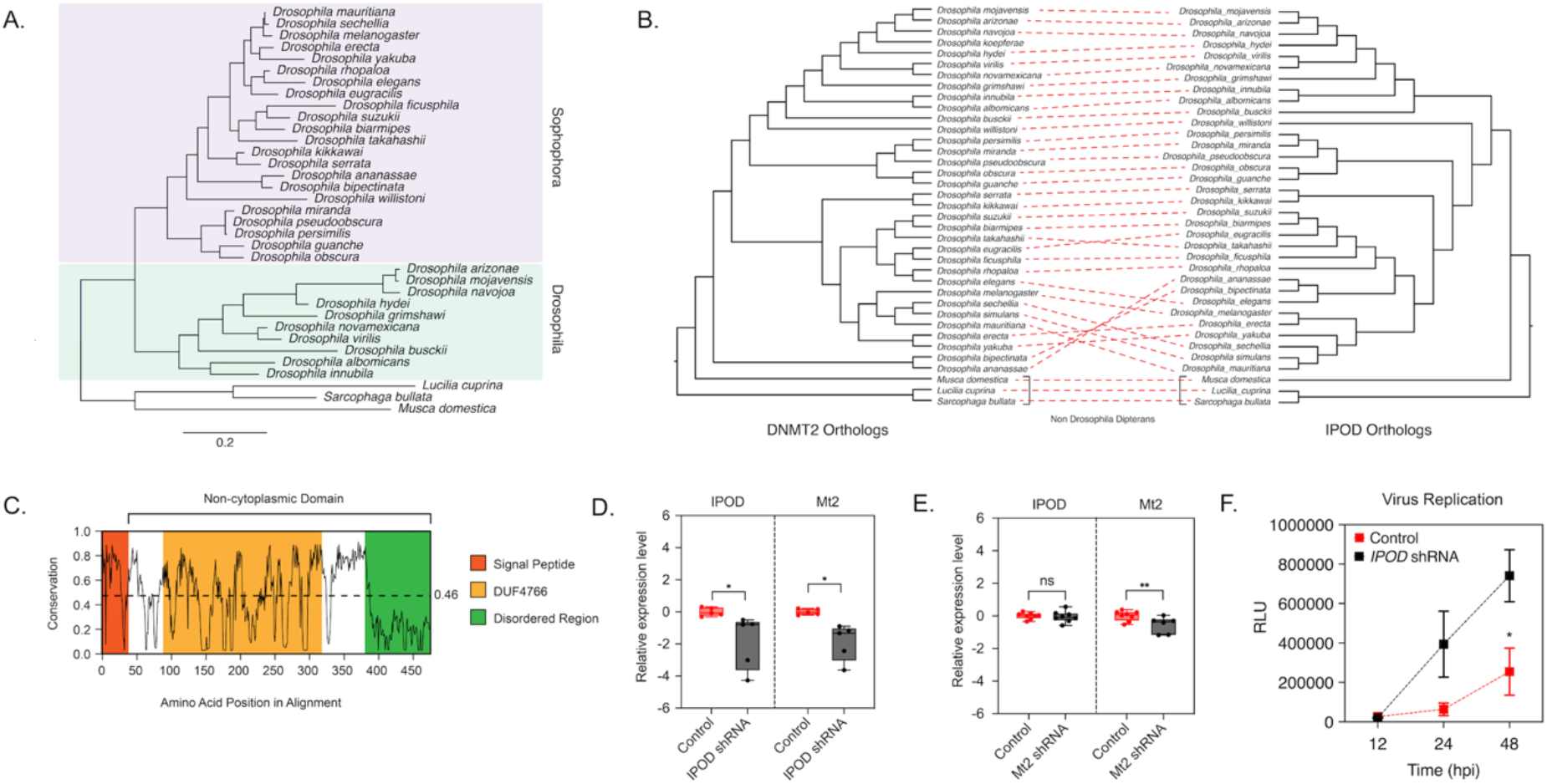
DNMT2 expression is regulated by IPOD in Drosophila species. (A) Maximum-likelihood (Deddouche, et al.) tree of the Interaction partner of DNMT2 (*IPOD*) gene present in multiple *Drosophila* species was constructed using RAxML using a multiple sequence alignment of *IPOD* nucleotide sequences. Sequence of the *IPOD* orthologs from *Lucilia cuprina, Musca domestica* and *Sarcophaga bullata* were used as outgroups. Scale bar represent branch lengths. (B) Inter-protein co-evolutionary analyses of DNMT2 and IPOD orthologs was performed using MirrorTree and TreeCmp software packages. Red, dashed lines connect the same Drosophila taxon. (C) Shannon conservation plot representing the degree of conservation (Y-axis) of IPOD orthologs present at every amino acid position (X-axis) across *Drosophilids*. Mean conservation score (0.46) across all amino acid positions is indicated by the horizontal dashed line. Colored boxes represent three InterPro domains identified across all IPOD orthologs in *Drosophilids*, including the N-terminal signal peptide (depicted in orange), followed by a C-terminal non-cytoplasmic domain (depicted in white) consisting of a conserved domain of unknown function (DUF4766, depicted in yellow) and a glycine-rich disordered region (depicted in green) present at the C-terminal end. (D,E) *IPOD* is an upstream regulator of *Mt2* expression in *Drosophila melanogaster*. (D) *IPOD* expression was knocked down in *Wolbachia w*Mel-colonized *Drosophila melanogaster* (TRiP line# 60092) by driving expression of a targeting short-hairpin RNA (shRNA) against the target mRNA. Relative expression of the target *IPOD* mRNA and *Mt2* mRNA was assessed via quantitative RT-PCR using total RNA derived from age-matched females. Siblings lacking the shRNA was used as the negative control. Two-tailed t-tests on log-transformed values; *IPOD*: p < 0.05, t = 3.678, df = 8.00, *Mt2*: p < 0.05, t = 2.454, df = 8.00. Error bars represent standard error of mean (SEM) of experimental replicates (n=5) (E) *Mt2* expression was knocked down in *Wolbachia w*Mel-colonized *Drosophila melanogaster* by driving expression of a targeting short-hairpin RNA (shRNA) against the target mRNA. Relative expression of the target *Mt2* mRNA and *IPOD* mRNA was assessed via quantitative RT-PCR using total RNA derived from age-matched females. Siblings lacking the shRNA was used as the negative control. Two-tailed t-tests on log-transformed values; *Mt2*: p < 0.01, t = 2.576, df = 12.00, *IPOD*: p = 0.717969, t = 0.3686, df = 14.00. Error bars represent standard error of mean (SEM) of experimental replicates (n=6-8). (F) Effect of *IPOD* knockdown on *Wolbachia-*mediated virus inhibition. Age-matched *Wolbachia-*colonized female flies either wild-type or expressing *IPOD*-targeting shRNA were intrathoracically injected with SINV-nLuc virus. At indicated times post infection (X-axis), flies were harvested and snap frozen prior to homogenization. Homogenized lysates were used to measure luciferase expression (RLU, Y-axis) which was subsequently used as a proxy to quantify virus replication. Two-way ANOVA of multivariate comparisons with Sidak’s post-hoc test; *IPOD* knockdown: p < 0.01, Time: p < 0.01. Error bars represent standard error of mean (SEM) of experimental replicates (n=3/time point). *p < 0.05, **p < 0.01, ns = not-significant.

In order to better understand IPOD’s cellular function, we performed a domain analyses using Pfam and InterPro. We identified a DUF4766 (PF15973) domain (Residues: 82 – 232) present in all orthologs, and InterPro suggested nearly 90% of the protein (Residues: 33 – 349) contains a non-cytoplasmic domain, with a smaller signal peptide domain (Residues: 1 – 32) present at the N-terminal end (Posterior Probability Score > 0.99) (Goll, et al. 2006; Ye, et al. 2018). Notably, we found nearly 28% (97/397) of the total protein length to be made of glycine residues, which are associated with a high degree of disordered structure. Indeed, an IUPred search predicted large stretches of intrinsically disordered regions along the entire length of the protein (Disorder Tendency Score > 0.5) indicating a potential role of IPOD in mediating complex protein-protein interactions (Dosztányi, et al. 2005; Dosztányi 2018). Taken together, these features are consistent with IPOD’s role as a nuclear protein with a potential role in transcriptional regulation. Interestingly, the distinct lack of a canonical DNA-binding domain within IPOD suggests that its ability to interact with other DNA-binding proteins is critical for its role in transcriptional regulation of DmDNMT2.

Given the absence of IPOD orthologs in the members of the Culicidae family e.g. *Aedes* mosquitoes, we hypothesized that IPOD regulates *Dm*DNMT2 (*Mt2*) expression, potentially replacing the role of miRNAs in the mosquito system. To validate IPOD’s role in *Dm*DNMT2 regulation, we used RNAi to knockdown IPOD (*IPOD*) expression *in vivo* in a transgenic fruit fly model by driving the expression of IPOD targeting short-hairpin RNA (shRNA) and measured relative mRNA levels of both *IPOD* and *Mt2* genes. We also measured these levels within the context of transgenic RNAi flies expressing shRNA against *Dm*DNMT2 to determine whether it affected levels of *IPOD* transcripts. Indeed, knocking down *IPOD* expression led to significantly reduced *Mt2* mRNA levels in flies expressing *IPOD*-targeting shRNA; Two-tailed t-tests on log-transformed values; *IPOD*: p < 0.05, t = 3.678, df = 8.00, *Mt2*: p < 0.05, t = 2.454, df = 8.00 (Figure 4D). Conversely, depleting *Dm*DNMT2 did not cause any significant change in *IPOD* mRNA levels, suggesting that *IPOD* likely functions upstream in the regulatory pathway; t-tests on log-transformed values, *IPOD, Mt2*: p < 0.01, t = 2.576, df = 12.00, *IPOD*: p = 0.717969, t = 0.3686, df = 14.00 (Figure 4E). Additionally, we wondered whether knockdown of IPOD affects virus inhibition within the context of a *Wolbachia-*colonized fly host. We reasoned that if IPOD is a positive regulator of *Dm*DNMT2 expression, its loss would lead to a subsequent reduction in *Dm*DNMT2 levels, thereby rescuing virus from *Wolbachia-*mediated inhibition, phenocopying our previous results (Bhattacharya, et al. 2017). Flies expressing *IPOD-*targeting shRNA were challenged with a SINV expressing a translationally fused luciferase reporter (SINV-nLuc) and virus replication at 12-, 24- and 48-hours post infection was measured by quantifying luciferase activity as a proxy for viral gene expression. Consistent with results obtained in our previous study, knockdown of *IPOD* in *Wolbachia-*colonized flies led to a significant increase in viral RNA, likely as a consequence of reduced *Dm*DNMT2 levels; Two-way ANOVA with Sidak’s post-hoc multiple comparisons test; *IPOD* knockdown: p < 0.01, Time: p < 0.01 (Figure 4F). This effect was independent of any change in endosymbiont titer (Unpaired Welch’s t-test: p = 0.4788, t = 0.7695, df = 4 (Supplementary Figure 5)). It should be noted that we have previously demonstrated *Wolbachia* titer does not change as a result of *Dm*DNMT2 *(Mt2)* knockdown in flies using the same experimental setup (Bhattacharya, et al. 2017). Taken together, these results support IPOD’s role in regulating *Dm*DNMT2 expression in the fruit fly. Furthermore, it notably demonstrates its importance in *Wolbachia*-mediated virus inhibition in terms of regulating *Dm*DNMT2 expression.

### Antiviral role of DNMT2 is host-dependent

Finally, we asked whether the antiviral role of *Dm*DNMT2 is a consequence of intrinsic features unique to this ortholog, or if this antiviral activity relies on specific interactions unique to its native host cell environment. To this end, we carried out heterologous expressions of *Dm*DNMT2 and *Aa*DNMT2 in their non-native *Aedes albopictus* and *Drosophila melanogaster* cells alongside their native counterparts and assessed their effect on virus. It should be noted, that ectopic expression of the non-native orthologs was carried out in the presence of the respective endogenous MTases. However, given the low levels of native DNMT2 expression in the cell, we reasoned that ectopic expression of the non-native ortholog should allow it to function as the dominant MTase variant.

Previous work has demonstrated that ectopic expression of *Dm*DNMT2 in *Drosophila melanogaster* derived JW18 cells causes reduction in infectious virus production, mirroring its antiviral role *in vivo* (Durdevic, Hanna, et al. 2013; Bhattacharya, et al. 2017). Altogether, *Dm*DNMT2 is able to restrict multiple viruses from at least four distinct RNA virus families highlighting a broad spectrum of antiviral activity. To determine whether this property is unique to *Dm*DNMT2 in the *Drosophila melanogaster* host, we expressed *Aa*DNMT2 in this host background and tested its effect on infectious virus production following challenge with the prototype alphavirus, SINV. *Drosophila melanogaster* derived JW18 cells (cleared of *Wolbachia* infection) were transfected with FLAG-tagged versions of *Dm*DNMT2 or *Aa*DNMT2 and were challenged with SINV at an MOI of 10 particles/cell approximately 72 hours post transfection. Cell supernatants were collected after 48 hours post infection and viral titers assayed on vertebrate baby hamster kidney fibroblast (BHK-21) cells using standard plaque assays. Consistent with our previous report, we saw a significant reduction in viral titer in cells expressing *Dm*DNMT2, compared to cells expressing the empty vector control. Notably, this result was phenocopied in cells expressing the non-native *Aa*DNMT2 ortholog; One-way ANOVA with Tukey’s post hoc test for multiple comparisons: Empty Vector vs *Dm*DNMT2: p = 0.0016, Empty Vector vs *Aa*DNMT2: p = 0.0017, *Dm*DNMT2 vs *Aa*DNMT2: p = 0.9971 (Figure 5B). We also assessed the effect of *Dm*DNMT2 and *Aa*DNMT2 expression on the per-particle infectivity of these progeny viruses, which is presented as the specific infectivity ratio of total infectious virus titer and total viral genome copies present in the cell supernatant (Bhattacharya, et al. 2020). In this case, expression of both DNMT2 orthologs were found to significantly reduce virion infectivity in cells compared to those expressing the empty vector; One-way ANOVA with Tukey’s post hoc test for multiple comparisons: Empty Vector vs *Dm*DNMT2: p = 0.0030, Empty Vector vs *Aa*DNMT2: p = 0.0066, *Dm*DNMT2 vs *Aa*DNMT2: p = 0.6951 (Figure 5C). These results indicate that like *Dm*DNMT2, *Aa*DNMT2’s MTase activity is antiviral within the context of the fruit fly, presumably through hypermethylation of the target viral and/or host RNAs (Figure 6).

**Figure 5.**
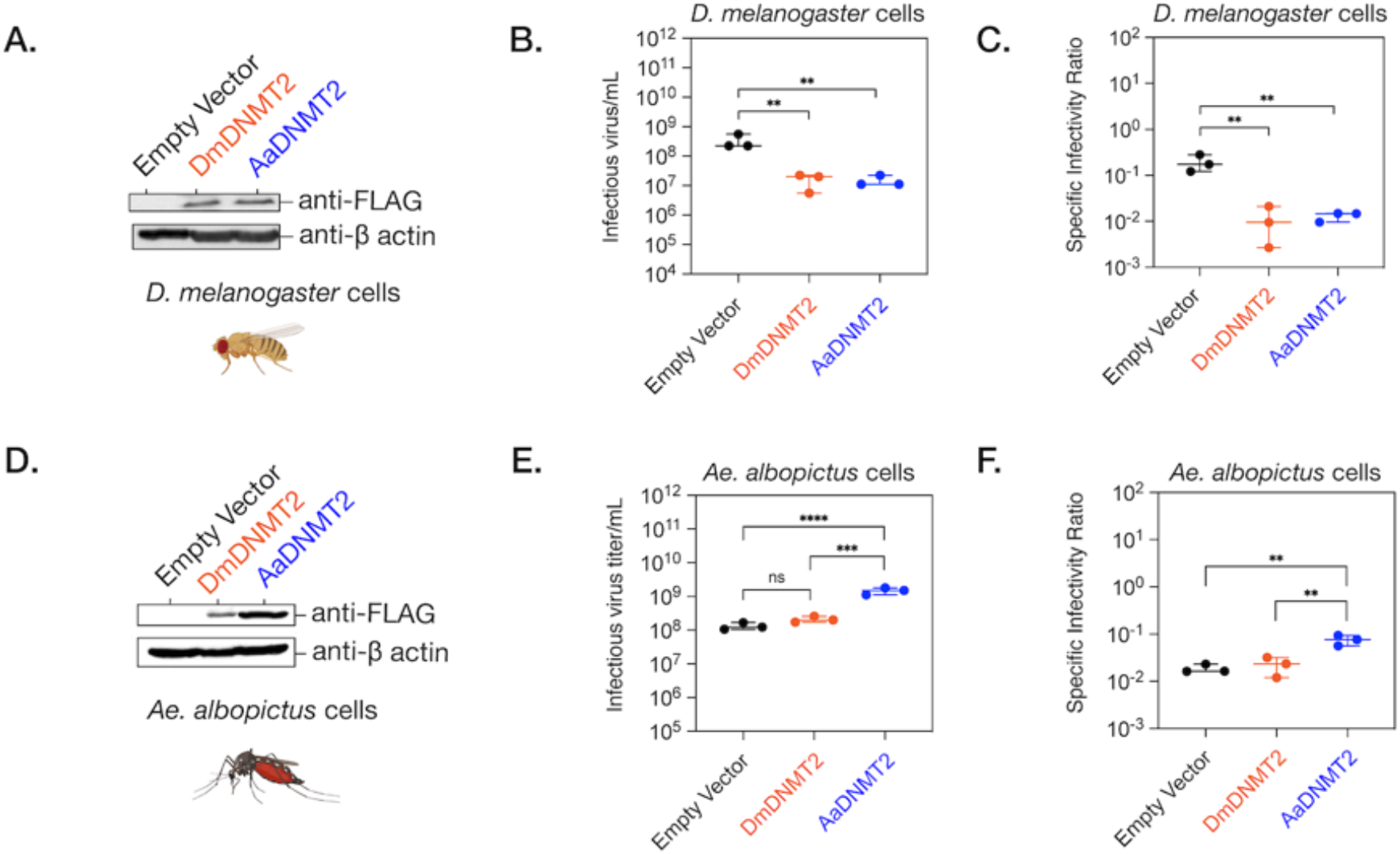
Effect of DNMT2 orthologs on virus replication is host-dependent. (A) *Drosophila melanogaster* derived JW18 cells (without *Wolbachia*) were transfected with plasmid constructs expressing epitope (FLAG) tagged versions of either the native fly (*Dm*DNMT2, depicted in orange) or the non-native mosquito (*Aa*DNMT2, depicted in blue) orthologs. Empty vector carrying only the FLAG-tag was used as a negative control (depicted in black). Protein expression was assessed 72 hours post transfection via Western Blot using antibodies against the FLAG-epitope. Cellular β-actin protein expression, probed using anti-β-actin antibody, was used as loading control. (B) 72 hours post transfection, JW18 cells expressing either the empty vector, the native DNMT2 (*Dm*DNMT2) or the non-native DNMT2 (*Aa*DNMT2) were challenged with SINV at MOI of 10 particles/cell. Cell supernatants were collected 48 hours post infection and infectious virus production was assessed via standard plaque assays on mammalian fibroblast BHK-21 cells. One-way ANOVA with Tukey’s post hoc test for multiple comparisons: Empty Vector vs *Dm*DNMT2: p = 0.0016, Empty Vector vs *Aa*DNMT2: p = 0.0017, *Dm*DNMT2 vs *Aa*DNMT2: p = 0.9971. Error bars represent standard error of the mean of 3 independent experiments. (C) Specific infectivity ratios of progeny SINV derived from JW18 cells either the empty vector, the native DNMT2 (*Dm*DNMT2) or the non-native DNMT2 (*Aa*DNMT2) was calculated as the ratio of infectious virus titer (infectious particles) and total viral genome copies (total virus particles) present in supernatants collected 72 hours post infection. One-way ANOVA with Tukey’s post hoc test for multiple comparisons: Empty Vector vs *Dm*DNMT2: p = 0.0030, Empty Vector vs *Aa*DNMT2: p = 0.0066, *Dm*DNMT2 vs *Aa*DNMT2: p = 0.6951. Error bars represent standard error of the mean of 3 independent experiments. (D) *Aedes albopictus* derived C636 cells (without *Wolbachia*) were transfected with plasmid constructs expressing epitope (FLAG) tagged versions of either the native fly (*Dm*DNMT2, depicted in orange) or the non-native mosquito (*Aa*DNMT2, depicted in blue) orthologs. Empty vector carrying only the FLAG-tag was used as a negative control (depicted in black). Protein expression was assessed 72 hours post transfection via Western Blot using antibodies against the FLAG-epitope. Cellular β-actin protein expression, probed using anti-β-actin antibody, was used as loading control. (E) 72 hours post transfection, *Aedes albopictus* derived C710 cells (colonized with *w*Stri Wolbachia strain) expressing either the empty vector, the native DNMT2 (*Dm*DNMT2) or the non-native DNMT2 (*Aa*DNMT2) were challenged with SINV at MOI of 10 particles/cell. Cell supernatants were collected 48 hours post infection and infectious virus production was assessed via standard plaque assays on mammalian fibroblast BHK-21 cells. One-way ANOVA with Tukey’s post hoc test for multiple comparisons: Empty Vector vs *Dm*DNMT2: p = 0.0937, Empty Vector vs *Aa*DNMT2: p < 0.0001, *Dm*DNMT2 vs *Aa*DNMT2: p = 0.0001. Error bars represent standard error of the mean of 3 independent experiments. (F) Specific infectivity ratios of progeny SINV derived from JW18 cells either the empty vector, the native DNMT2 (*Dm*DNMT2) or the non-native DNMT2 (*Aa*DNMT2) was calculated as the ratio of infectious virus titer (infectious particles) and total viral genome copies (total virus particles) present in supernatants collected 72 hours post infection. One-way ANOVA with Tukey’s post hoc test for multiple comparisons: Empty Vector vs *Dm*DNMT2: p = 0.8969, Empty Vector vs *Aa*DNMT2: p = 0.0060, *Dm*DNMT2 vs *Aa*DNMT2: p = 0.0095. Error bars represent standard error of mean of 3 independent experiments. ****p < 0.0001, ***p < 0.001, **p < 0.01, ns = not-significant.

**Figure 6.**
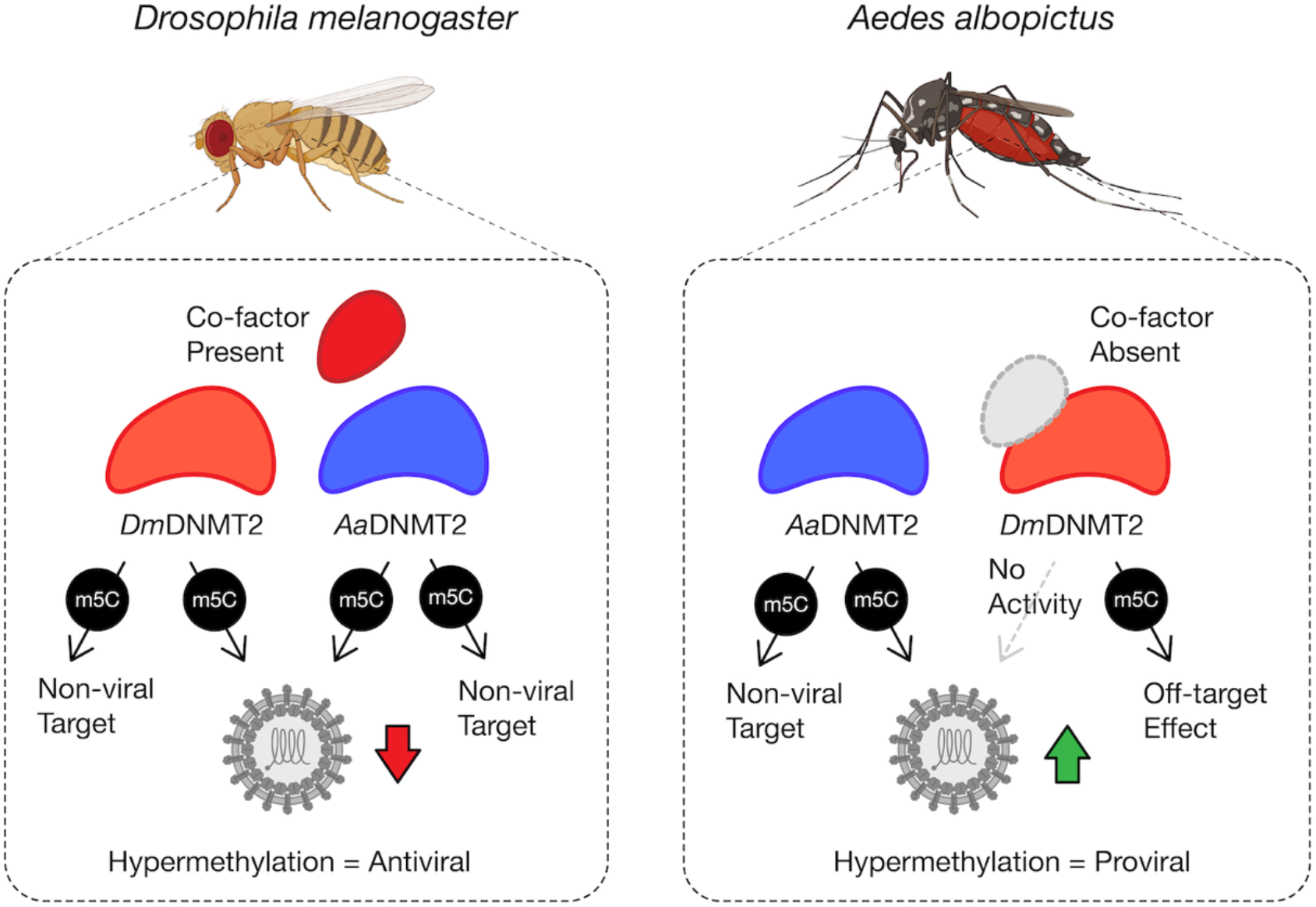
Model schematic of *Dm*DNMT2 and *Aa*DNMT2 activity. Heterologous expression of either *Dm*DNMT2 or *Aa*DNMT2 in *Drosophila melanogaster* derived JW18 cells leads to virus inhibition, likely as a consequence of hypermethylation of a viral and/or host target. In this case, *Dm*DNMT2 function is potentially aided by the presence of an unidentified co-factor. Heterologous expression of *Aa*DNMT2 in *Wolbachia-* colonized *Aedes albopictus* cells leads to the rescue of virus inhibition, likely due to hypermethylation of a viral and/or host target. In contrast, *Dm*DNMT2 expression in these cells has no observable effect on virus replication suggesting either a loss in MTase activity or potential off-target effects. This result could be due to the absence of *Dm*DNMT2’s cognate interaction partner(s) or co-factor(s) that are unique to Drosophila and are thus absent in *Aedes albopictus* cells.

At this point we should emphasize that *Aa*DNMT2 has been shown to be proviral in the *Aedes* context. While *Wolbachia* upregulates *Dm*DNMT2, leading to virus inhibition, colonization by *Wolbachia* in *Aedes* backgrounds reduces *Aa*DNMT2 expression, leading to RNA virus restriction (Zhang *et al*., 2013). At the same time, *Aa*DNMT2 expression is induced in the presence of virus alone, implying a proviral role that is lost in the presence of *Wolbachia*. We therefore reasoned that ectopic expression of *Aa*DNMT2 should rescue virus from *Wolbachia-*mediated inhibition in *Aedes* mosquito cells. We wondered, therefore, if heterologous expression of *Dm*DNMT2 in the *Aedes* cellular context would result in a proviral phenotype. *Aedes albopictus* (C710) derived cells (colonized with the *w*Stri *Wolbachia* strain) were transfected with FLAG-tagged versions of *Dm*DNMT2 or *Aa*DNMT2 and were challenged with SINV at an MOI of 10 particles/cell approximately 72 hours post transfection. As before, cell supernatants were collected after 48 hours post infection and viral titers assayed on vertebrate baby hamster kidney fibroblast (BHK-21) cells using standard plaque assays. In line with our hypotheses, expression of *Aa*DNMT2 in these cells was associated with a significant increase in SINV titer. However, we did not find any significant changes in virus titer from cells expressing the non-native *Dm*DNMT2 ortholog; One-way ANOVA with Tukey’s post hoc test for multiple comparisons: Empty Vector vs *Dm*DNMT2: p = 0.0937, Empty Vector vs *Aa*DNMT2: p < 0.0001, *Dm*DNMT2 vs *Aa*DNMT2: p = 0.0001 (Figure 5E). We also observed a similar trend after measuring the per-particle infectivity of these progeny viruses, with an increase in virion infectivity upon expression of *Aa*DNMT2 but not *Dm*DNMT2; One-way ANOVA with Tukey’s post hoc test for multiple comparisons: Empty Vector vs *Dm*DNMT2: p = 0.8969, Empty Vector vs *Aa*DNMT2: p = 0.0060, *Dm*DNMT2 vs *Aa*DNMT2: p = 0.0095 (Figure 5F). Additionally, we assessed the effect of heterologous *Dm*DNMT2 on viral RNA levels in the cell based on previous reports that demonstrated the ability of *Aa*DNMT2 to rescue virus replication in the presence of *Wolbachia* (Zhang, et al. 2013). Consistent with previous findings, expression of *Aa*DNMT2 significantly improved SINV RNA levels in cells. However, heterologous expression of *Dm*DNMT2 did not have any effect on SINV RNA levels; One-way ANOVA with Tukey’s post hoc test for multiple comparisons: SINV RNA, Empty Vector vs *Dm*DNMT2: p = 0.7875, Empty Vector vs *Aa*DNMT2: p < 0.05, *Dm*DNMT2 vs *Aa*DNMT2: p < 0.05 (Supplementary Figure 5A). Finally, we quantified *Wolbachia* gene expression across these conditions to ensure that any changes in virus fitness was not caused due to changes in endosymbiont titer. We did not find any evidence of either *Aa*DNMT2 or *Dm*DNMT2 expression to have any effect on *Wolbachia wsp* gene expression; Empty Vector vs *Dm*DNMT2: p = 0.4121, Empty Vector vs *Aa*DNMT2: p = 0.5639, *Dm*DNMT2 vs *Aa*DNMT2: p = 0.9523 (Supplementary Figure 5B).

Altogether these results suggest that heterologous expression of *Dm*DNMT2 in mosquito cells has no effect on virus fitness. We hypothesize that the cellular context of expression determines the interaction between host, DNMT2 ortholog, and virus. Given the role of Drosophila-specific host factor IPOD in regulating *Dm*DNMT2 antiviral function (Figure 4), and the detected adaptive changes on DNMT2’s surface, we speculate that this lack of *Dm*DNMT2 activity in mosquito cells (Figure 5E,F) might occur due to the lack of one or more interaction partners or co-factors.

## Discussion

Here we present a functional analysis of adaptive evolution of DNMT2 in Dipteran insects that adds support to recent reports describing its role in host innate immunity (Durdevic, Hanna, et al. 2013; Durdevic and Schaefer 2013; Zhang, et al. 2013; Bhattacharya, et al. 2017; Baradaran, et al. 2019; Claudio-Piedras, et al. 2019). The biological function of DNMT2 remains unexplored in a vast majority of arthropods. Where it has been studied, for example in *Drosophila melanogaster*, loss of function of DNMT2 is not associated with any severe developmental issues or lethality (Goll, et al. 2006). Additionally, DNMT2-only insects like fruit flies and other holometabolous insects exhibit very low to no CpG methylation across their genome, in line with DNMT2’s lack of DNA MTase activity (Lewis, et al. 2020). Recent studies suggest that DNMT2 is a part of the cellular stress response that also acts against external stressors like pathogen challenges. Indeed, *Dm*DNMT2 confers protection against a wide range of RNA viruses and bacteria like *Acetobacter tropocalis, Lactobacillus fructivorans* and *Acetobacter pomorum* (Durdevic, Hanna, et al. 2013; Bhattacharya, et al. 2017). Similarly, the DNMT2 ortholog in *Helicoverpa armigera* (Order: Lepidoptera) has been shown to confer protection against systemic infections by *Bacillus thuringiensis* and *Serratia marcescens* (Baradaran, et al. 2019). However, there are instances where DNMT2 regulates how well certain pathogens colonize the host in a manner that is seemingly beneficial to the former. Examples of this can be found among members of the Culicidae family (Zhang, et al. 2013; Claudio-Piedras, et al. 2019). In each of these cases, expression of DNMT2 is elevated following an infectious bloodmeal containing either the parasite *Plasmodium berghei (Anopheles albimanus)* or DENV *(Aedes aegypti)* (Zhang, et al. 2013; Claudio-Piedras, et al. 2019). Notably, pharmacological inhibition or miRNA-mediated knockdown of DNMT2 in these species correlates with reduced host susceptibility to infection. However, it is clear from these examples that Drosophilidae and Culicidae DNMT2 plays an important role in shaping the host immune response to a wide range of pathogens, notably RNA viruses (Durdevic, Hanna, et al. 2013; Durdevic and Schaefer 2013; Zhang, et al. 2013; Bhattacharya, et al. 2017; Baradaran, et al. 2019; Claudio-Piedras, et al. 2019).

### Elucidating the molecular evolution of DNMT2

Signatures of positive selection are often a hallmark of genes involved in host immunity (Moutinho, et al. 2019). To determine whether DNMT2 itself has been subjected to such selection, we carried out CodeML analyses of DNMT2 orthologs from Dipteran insects, with an increased focus on members of the Culicidae and Drosophilidae families based on their roles in host immunity (Yang 2007). In previous studies, we and others have described the role of *Dm*DNMT2 and *Aedes* DNMT2 orthologs in *Wolbachia-*mediated inhibition of RNA viruses (Zhang, et al. 2013; Bhattacharya, et al. 2017). DNMT2 is thought to interact with the viral RNA in the cytoplasm and influence virus replication in a manner that is dependent on their catalytic activity (Durdevic et al., 2013). Furthermore, overexpression or loss-of-function of *Dm*DNMT2 caused a corresponding increase and reduction in virus restriction, while the reverse phenotype is observed for *Aa*DNMT2 (Bhattacharya et al., 2017; Zhang et al., 2013). Indeed, overexpression of *Aa*DNMT2 caused a corresponding increase in virus replication, respectively, indicating a pro-viral role for this ortholog (Zhang et al., 2013). Consistent with known roles of *Dm*DNMT2 and *Aa*DNMT2 in virus regulation, in this study we found several instances of positive selection along ancestral and more recent lineages leading to these species, identifying several potential codon sites within each ortholog having experienced positive selection (Figure 2, Tables 1-2). Notably, our results regarding the presence of positive selection in the lineage ancestral to Sophophora subgenus are consistent with a recent study by Vieira et.al., and 3 specific residues (87T, 261L, 320K) were identified in both analyses (Vieira, et al. 2018). Physiochemical properties and location of these amino acid residues on the 3D structure of *Dm*DNMT2 and *Aa*DNMT2 indicate that these adaptive changes occur in four major regions of the protein (Figure 2A). Collectively, these changes might influence catalytic function and inter-molecular interactions with other accessory proteins and/or nucleic acid substrates. Further work, using site-directed mutagenesis of these sites, is required to validate the importance of these residues on the ability of these DNMT2 orthologs to regulate virus infection. Notably, our CodeML analyses did not find any evidence of positive selection along lineages leading to *Aedes* DNMT2 since their divergence with *Anopheles* (Figure 1A). This is in contrast with the antiviral role of *Dm*DNMT2, which could explain the presence of positive selection along this lineage. However, several sites identified in the ancestral Culicidae lineage as well as related *Anopheles* genera were found to occur within *Aa*DNMT2 (Figure 1A). Furthermore, heterologous expression of this ortholog in fly cells were able to restrict infectious virus production as well as the native *Dm*DNMT2 ortholog, indicating that the outcome is host-dependent (Figure 6). Collectively, our results suggest that several Dipteran DNMT2 orthologs may have evolved to function at the interface of host-pathogen interactions, contributing to its antiviral role in fruit flies and possibly other members of the Drosophila genus. Indeed, based on overall positive selection and complete conservation of these codon sites among Drosophila/Sophophora, it is conceivable that these DNMT2 orthologs confer similar antiviral effects in their respective host backgrounds (Figure 2, Tables 1-2). Given the lack of genetic tractability in these Drosophila species, heterologous expression of these DNMT2 orthologs in a tractable *Drosophila melanogaster* background can be used to determine their restriction properties.

### Delineating differences between DNMT2 regulation in fruit fly and mosquitoes

In addition to the presence or absence of positive selection, we identified two distinct differences in the overall protein sequence between Drosophilidae and Culicidae DNMT2. The first being an extended (7-47 aa long), unstructured N-terminal end present in all DNMT2 orthologs within Culicidae species. The other difference lies in the target recognition domain, which is extended (7-11 aa long) in Drosophilidae DNMT2 and is predicted to interact with the nucleic acid substrate based on past simulation studies using mammalian DNMT1 (Ye, et al. 2018). These differences also give rise to altered surface charge distribution between *Dm*DNMT2 and *Aa*DNMT2, further signifying potential differences in inter-molecular associations and/or target specificity between these orthologs. These differences could represent unique modes of regulation between the two orthologs, a case strengthened by our results regarding the role of the *Drosophila melanogaster* protein IPOD in *Dm*DNMT2 regulation. IPOD is present within all members of the Drosophila genus, but absent in Culicidae species (Supplementary Figure 4). Notably, previous *in vivo* and *in vitro* analyses indicate that IPOD binds to the N-terminal end of DmDNMT2 (Kunert 2005). Primary amino acid sequence composition of IPOD also indicates a vast portion of this protein to be intrinsically unstructured, suggesting a great degree of conformational flexibility that might allow extensive protein-protein interactions. Previous work has also suggested IPOD-mediated regulation of *Dm*DNMT2 expression. Through *in vivo* loss-of-function analyses, we show that IPOD is indeed an upstream regulator of *Dm*DNMT2 expression. Given that the entirety of IPOD is made up of an N-terminal signal peptidase and a C-terminal non-cytoplasmic domain, it is likely that it regulates *Dm*DNMT2 transcription in the nucleus. Finally, demonstrating its functional role in *Dm*DNMT2 regulation, we show that loss of IPOD in flies colonized with *Wolbachia* phenocopy *Wolbachia-*colonized *Dm*DNMT2 loss-of-function mutants (Bhattacharya, et al. 2017). Consistent with our previous reports, this loss in virus inhibition occurs without any changes in endosymbiont titer. The role of IPOD as a cognate DNMT2 regulator and interaction partner is further supported by our observation that the phylogenies of Drosophila DNMT2 and IPOD orthologs mirror one another to a significant degree, suggesting a co-evolving relationship between these two proteins.

The mechanism of Culicidae DNMT2 regulation is less well defined, but likely varies between different mosquito genera. A recent study by Claudio-Piedras et al. suggest that DNMT2 in *Anopheles albimanus* is under the control of the NF-κB family of transcription factors (Claudio-Piedras, et al. 2019). This is in contrast to Aedes mosquitoes, where expression of DNMT2 is under the control of a conserved miRNA *aae-miR-2940* (miRBase Accession: MI0013489) (Zhang, et al. 2013). However, like the miRNA itself, its target mRNA sequence is unique to Aedes DNMT2 and are absent from ortholog transcripts from other Culicidae species and most notably, from Drosophila DNMT2 (Supplementary Figure 6A). Still absence of this particular miRNA target does not imply that *Dm*DNMT2 is not under the control of any miRNAs. *In silico* miRNA prediction with *Dm*DNMT2 (FBtr0110911) as a target query using miRanda predicts one highly conserved host miRNA, dme-miR-283 (miRBase Accession ID: MI0000368), with the potential of targeting the 3’ untranslated region (3’UTR) of the *Dm*DNMT2 gene. Incidentally, dme-miR-283 is among the top ten most upregulated miRNAs in fly cells following alphavirus (Semilki Forest Virus, SFV) infection, both in the presence and absence of *Wolbachia* (Rainey, et al. 2016). Assuming that dme-miR-283 downregulates *Dm*DNMT2 expression, the modENCODE RNA-seq treatments dataset and our previous observations indicate these results are in line with the SINV-responsive expression pattern of this miRNA and its target in adult flies (Bhattacharya, et al. 2017). It should also be noted, that while we found a single miRNA targeting *Dm*DNMT2, miRanda and TargetScanFly v7.2 identified a set of three conserved Drosophila miRNAs targeting the 3’UTR region of multiple Drosophila IPOD orthologs (FBgn0030187). A subset of these miRNAs has been previously associated with regulating host innate immunity and antimicrobial responses (Supplementary Figure 6B) (Li, et al. 2017). Further work is necessary to experimentally validate the role of these miRNAs in regulating expression of their predicted targets.

### Influence of host backgrounds on DNMT2 antiviral activity

Finally, through heterologous expression of *Dm*DNMT2 and *Aa*DNMT2 in their non-native host backgrounds, we show that the antiviral activity is not unique to *Dm*DNMT2 but is rather a consequence of the host *Drosophila melanogaster* background, as its effect on SINV is phenocopied by heterologous *Aa*DNMT2 expression in the same cells, leading to a loss in infectious virus production as well as per-particle infectivity. This suggests that sequence or structural features that are unique to *Dm*DNMT2 are not responsible for its antiviral activity in fly cells. However, these features do indicate the requirement for specific inter-molecular interactions that is required for proper *Dm*DNMT2 function and specificity. This is supported by our observation that expression of *Dm*DNMT2 in *Aedes albopictus* mosquito cells has no effect on SINV, either antiviral or proviral, in contrast to the native *Aa*DNMT2 expression which leads to virus “rescue” from *Wolbachia-*mediated inhibition. We postulate that this complete lack of *Dm*DNMT2 activity and/or specificity in this host *(Aedes albopictus)* background could be due to the absence of one or more *Dm*DNMT2 “co-factors” that are specific to Drosophila i.e. IPOD (Figure 6).

Our observations regarding *Aa*DNMT2’s ability to function as an antiviral in fly cells suggests that any selection within Drosophila that differs from Aedes may also be due to other adaptations. Still, the sites identified to be under positive selection may contribute to *Dm*DNMT2’s potency as an antiviral. Further work is required to determine if *Dm*DNMT2 variants carrying the replaced ancestral codons are less efficient at inhibiting viruses native to Drosophila, as they likely represent the source of this selection.

Since the exact mechanism of DNMT2’s antiviral role remains undefined, it is possible that these adaptations allow for functional differences of this MTase against specific viruses, host conditions or both. Notably, the viruses used in this study are alphaviruses, which are native to the *Aedes* host. The antiviral activity of both MTase orthologs against these viruses in fly cells could therefore also be due to fundamental differences in the host response to potential hypermethylation of viral and host RNA species. Indeed, while such modifications may be favorable or even necessary for alphavirus replication in the native mosquito, it might allow for virus recognition and clearance in the fly background. Further studies are required using native virus-host-MTase ortholog combinations to explore these possibilities. At the same time, based on our current experimental setup we cannot rule out the possibility that basal-level expression of the endogenous MTase has an effect on the outcomes of our heterologous-expression experiments. Further work is required to determine if heterologous expression of *Aa*DNMT2 can complement the absence of the native-*Dm*DNMT2 null fly cells, and vice versa, with regards to virus restriction or rescue, respectively.

## Materials and Methods

### Insect and Mammalian Cell Culture

JW18 *Drosophila melanogaster* cells with and without *Wolbachia* (strain *w*Mel) were grown at 24 °C in Shields and Sang M3 insect media (Sigma-Aldrich) supplemented with 10% heat-inactivated fetal bovine serum (Gibco), 1% each of L-Glutamine (Corning), non-essential amino acids (Corning) and penicillin-streptomycin-antimycotic (Corning). Baby hamster kidney fibroblast (BHK-21) cells were grown at 37 °C under 5% CO_2_ in 1X Minimal Essential Medium (Corning) supplemented with 10% heat-inactivated fetal bovine serum (Corning), 1% each of L-Glutamine (Corning), non-essential amino acids (Corning) and penicillin-streptomycin-antimycotic (Corning).

### Fly husbandry, genetic crosses and virus injections

The following stocks were obtained from the Bloomington *Drosophila* Stock Center (BDSC) located at Indiana University Bloomington (http://flystocks.bio.indiana.edu/). *Wolbachia*-infected RNAi mutant stock 60092 (y[1] sc[*] v[1] sev[21]; P{y[+t7.7] v[+t1.8]=TRiP.HMC05086}attP40) was used for shRNA-mediated knock-down of IPOD gene expression by driving dsRNA expression using previously described Act5C-Gal4 driver males (y^1^ w^*^; P{w[Act5C-GAL4}17bFO1/TM6B, Tb^1^). The homozygous TRiP mutant adult females colonized with *Wolbachia* were crossed to uninfected w; Sco/Cyo males. Virgin progeny females carrying the inducible shRNA construct were collected and age-matched (2-5 days old) before being crossed to the aforementioned Act5C-Gal4 driver males. As per our previous study, *Wolbachia*-infected TRiP mutant stock 42906 (y1 sc* v1; P {TRiP.HMS02599} attP40) was used to achieve knock-down of *Mt2* gene expression by driving dsRNA expression using the aforementioned Act5C-Gal4 driver males. All fly stocks were maintained on standard cornmeal-agar medium diet supplemented with an antibiotic cocktail which comprises penicillin and streptomycin (P/S) at 25°C on a 24-hour light/dark cycle. In order to establish a systemic virus infection *in vivo*, flies were anesthetized with CO_2_ and injected intrathoracically with 50nL of approximately 10^10^ PFU/mL of purified Sindbis virus (SINV-nLuc) or sterile saline solution (1XPBS) using a nano-injector (Drummond Scientific). Flies were collected two days post-infection, snap frozen in liquid N_2_ and stored at -80 °C for downstream processing. Samples for quantitative PCR and quantitative RT-PCR were homogenized in TRiZOL reagent (Sigma Aldrich) and further processed for nucleic acid extractions using manufacturer’s protocols.

### DNMT2 overexpression in insect cells

Expression vectors containing *Drosophila melanogaster* and *Aedes albopictus* DNMT2 orthologs used here were designed in the following manner; *Aedes albopictus AMt2* coding region was subcloned into PCR 2.1 TOPO vector (Invitrogen) by PCR amplification of cDNA generated using reverse transcribed from total cellular RNA isolated from C636 *Aedes albopictus* cells using Protoscript II RT (NEB) and oligo-dT primers (IDT). Coding region was validated via sequencing before cloned into the pAFW expression vector (1111) (Gateway Vector Resources, DGRC), downstream of and in-frame with the 3X FLAG tag using the native restriction sites AgeI and NheI (NEB). Expression of both FLAG-tagged AaDNMT2 in mosquito cells was confirmed using qRT-PCR and Western Blots using an anti-FLAG monoclonal antibody (SAB4301135 - Sigma-Aldrich) (Fig 4A). Catalytic MTase mutant of *AMt2* (*AMt2*-C78G), was generated via site-directed mutagenesis (NEB, Q5 Site-Directed Mutagenesis Kit). using primers listed in the primer table (Table S1). *Drosophila Mt2* (FBgn0028707) cDNA clone (GM14972), obtained from DGRC (https://dgrc.bio.indiana.edu/) was cloned into the pAFW expression vector (1111) with an engineered SaII site (Gateway Vector Resources, DGRC) downstream of and in-frame with the 3X FLAG tag using Gibson assembly (HiFi DNA assembly mix, NEB). Expression of FLAG-tagged DNMT2 in fly cells was confirmed using qRT-PCR and Western Blots using an anti-FLAG monoclonal antibody (SAB4301135 - Sigma-Aldrich). Catalytically inactive *Mt2* (*Mt2* C78A) variant was generated via site-directed mutagenesis (NEB, Q5 Site-Directed Mutagenesis Kit) using primers listed in the primer table (Supplementary Table 1). JW18 fly cells were transfected with expression constructs using Lipofectamine LTX supplemented with Plus reagent (Invitrogen) by following manufacturer’s protocol. Protein expression was assessed 72 hours post transfection via Western Blot using a monoclonal antibody against the FLAG epitope (Sigma) (Figure 5A, D). For every western blot experiment, monoclonal anti-β-actin antibody was used to probe cellular β-actin levels, which was used as loading control.

### Virus infection in cells

Viral titers were determined using standard plaque assays on baby hamster kidney fibroblast (BHK-21) cells. Cells were fixed 48 hours post infection using 10% (v/v) formaldehyde and stained with crystal violet to visualize plaques. Virus particles were determined by quantifying viral genome copies via quantitative RT-PCR using primers targeting the SINV E1 gene (See Supplementary Table 1 for primer details) and standard curves made from linearized infectious clone sequences containing full-length SINV genome. Primer efficiencies for this primer set was determined in our previous study (Bhattacharya, et al. 2020).

### Real-time quantitative PCR and RT-PCR analyses

Total DNA and RNA were extracted from samples using TRiZOL reagent (Sigma Aldrich) according to manufacturer’s protocols. Synthesis of complementary DNA (cDNA) was carried out using MMuLV Reverse Transcriptase (NEB) and random hexamer primers (Integrated DNA Technologies). Negative (no RT or no gDNA or cDNA synthesized from mock infected cell supernatants) controls were used for each target per reaction. Quantitative PCR or RT-PCR analyses were performed using Brilliant III SYBR green QPCR master mix (Bioline) with gene-specific primers on a Applied Bioscience StepOnePlus qPCR machine (Life Technologies). All primer sets were designed based on information present in existing literature [14,17,20]. Query gene expression levels were normalized to the endogenous 18S rRNA expression using the delta-delta comparative threshold method (ΔΔCT) (Supplementary Table 1).

### Phylogenetic Analyses

Maximum likelihood trees were constructed using RAxML using the Le-Gascuel (Bhattacharya, et al.) amino acid substitution model with 100 bootstrap replicates (Stamatakis 2014). Multiple sequence alignments were generated using Clustal Omega. Final trees were visualized using FigTree v1.4.4.

### CodeML analyses

Tree topologies were obtained using RAxML with aligned codon-based nucleotide sequences. The “-m GTRGAMMA” model was used with rapid bootstrap analysis and search for the best tree (option: -f a). The codeML null and alternative branch-site models were run for each individual branch in the tree as foreground independently (Yang 2007). In the alternative model, the branch site model allows a class of sites in the foreground branch to have a dN/dS > 1. In the text we refer generally to dN/dS as ω and to the dN/dS > 1 class as ω2. Convergence issues were addressed by rerunning analyses with different values for Small_Diff. Signs of convergence issues include: 1) lnL values worse than the M1a NearlyNeutral site model, 2) the first two site classes having proportions of zero, 3) the null model having better lnL than the alternative model, 4) in the alternative model, lnL values worse than expected given estimated site posterior probabilities.

### Inter-protein co-evolution analyses

Co-evolution of Drosophila DNMT2 and IPOD orthologs was performed using multiple sequence alignments using the MirrorTree Server (Ochoa and Pazos 2010). Robinsin-Foulds distances was calculated to measure the dissimilarity between the topologies of unrooted IPOD and DNMT2 phylogenetic trees using the Visual TreeCmp webserver (Bogdanowicz, et al. 2012; Bogdanowicz and Giaro 2017). The following optional parameters were selected for Matching Split and Weighted Robinsin Foulds aka. RFWeighted (0.5) and RF (0.5) analyses: Normalized distances, Prune trees (to allow for partially overlapping sets of taxa) and Zero weights allowed.

### *In silico* miRNA prediction

Prediction of miRNAs targeting Drosophila Mt2 (FBgn0028707) and IPOD (FBgn0030187) was carried out using two independent miRNA prediction servers, TargetScanFly v7.2 and microrna.org (Agarwal, et al. 2018; Kozomara, et al. 2019). The latter combines miRanda target prediction with an additional mirSVR target downregulation likelihood score (Betel, et al. 2010). Accession numbers of miRNAs predicted in this study was obtained from miRBase.

### Protein Conservation

Protein conservation was determined with the Protein Residue Conservation Prediction tool (http://compbio.cs.princeton.edu/conservation/index.html;) (Dosztányi, et al. 2005; Dosztányi 2018). Multiple sequence alignment of amino acid sequences carried out using Clustal Omega were used as input, while Shannon entropy scores were selected as output, alongside a window size of zero, and sequence weighting set to “false.” Conservation was subsequently plotted using GraphPad Prism 8. DNMT2 motif regions were defined as per described in previous studies (Falckenhayn, et al. 2016). For IPOD, domains were defined based on Pfam and InterPro domain prediction results obtained using *Drosophila melanogaster* IPOD as an input query (Capra and Singh 2007; Kelley, et al. 2015).

### Homology Modelling of DNMT2 orthologs

Template-based comparative modeling of DNMT2 orthologs from *Drosophila melanogaster, Aedes albopictus* and *Anopheles gambiae* was performed using the intensive modelling approach in Protein Homology/Analogy Recognition Engine 2 (Phyre2) [28]. Protein structures were visualized using PyMOL (The PyMOL Molecular Graphics System, Version 1.2r3pre. Schrödinger, LLC).

### Inter-protein co-evolution analyses

Co-evolution of Drosophila DNMT2 and IPOD orthologs was performed using multiple sequence alignments using the MirrorTree Server (Ochoa and Pazos 2010). Robinsin-Foulds distances was calculated to measure the dissimilarity between the topologies of unrooted IPOD and DNMT2 phylogenetic trees using the Visual TreeCmp webserver (Bogdanowicz, et al. 2012; Bogdanowicz and Giaro 2017). The following optional parameters were selected for Weighted Robinsin Foulds, RFWeighted (0.5) and RF (0.5) analyses: Normalized distances, Prune trees and Zero weights allowed.

### Statistical analyses of experimental data

All statistical tests were conducted using GraphPad Prism 8 (GraphPad Software Inc., San Diego, CA). Details of statistical tests for each experiment can be found in the results section and the associated figure legends.

### Graphics

Graphical assets made in BioRender - biorender.com.

## Supporting information

Accession numbers used in the study

## Acknowledgements

We would like to thank Dr. Suchetana Mukhopadhyay for her critical and helpful suggestions regarding interpretations of data included in this manuscript. We would like to thank members of the Hardy, Newton, Danthi, Mukhopadhyay and Patton labs for their critical input and thoughtful discussions. This project was supported by funding from NIH NIAID to ILGN (R01 AI144430) and to ILGN and RWH (R21 AI153785)

## Data availability

All sequences used in this manuscript are publicly available and accession numbers provided in Supplementary Materials.

## SUPPLEMENTARY FIGURES

**Supplementary Figure 1.**
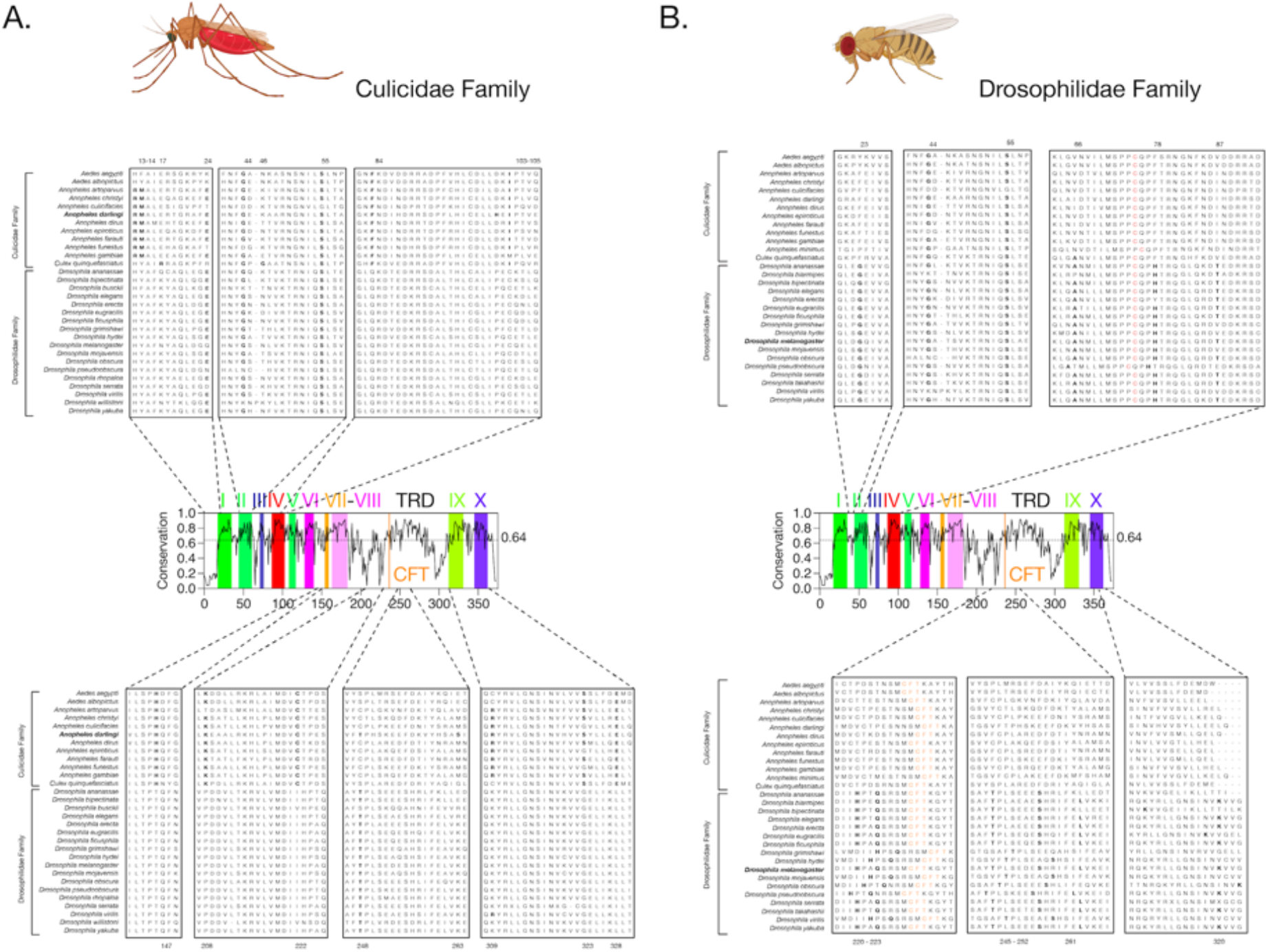
Amino-acid positions in Culicidae and Drosophila DNMT2 under positive selection. The global degree of primary sequence conservation (Shannon Conservation Score, Y-axis) of DNMT2 orthologs present at every amino acid position (X-axis) across *Drosophila* and *Culicidae* species. Colored boxes represent a total of ten canonical sequence motifs conserved within eukaryotic DNMT2, in addition to the DNA/RNA binding CFT motif located in the otherwise variable DNMT2 target recognition domain (TRD). Horizontal dotted line and the associated number on the left represent the mean Shannon conservation score averaged across all amino acid positions. Conservation of the amino acid positions in DNMT2 orthologs across *Drosophilids* and *Culicidae* species. Multiple sequence alignment of DNMT2 amino acid sequences was performed using U-Gene. Potential evidence of positive selection was identified along several branches (Figure 1, Table 1). Associated amino acid positions with high posterior probability values (> 95%) were considered as sites under selection (see Tables 1 and 2 for details) and are represented in bold letters. Catalytic Cysteine (Cys, C) residue within Motif IV is represented in red, bold letters. DNA/RNA binding CFT Motif is represented in orange, bold letters.

**Supplementary Figure 2.**
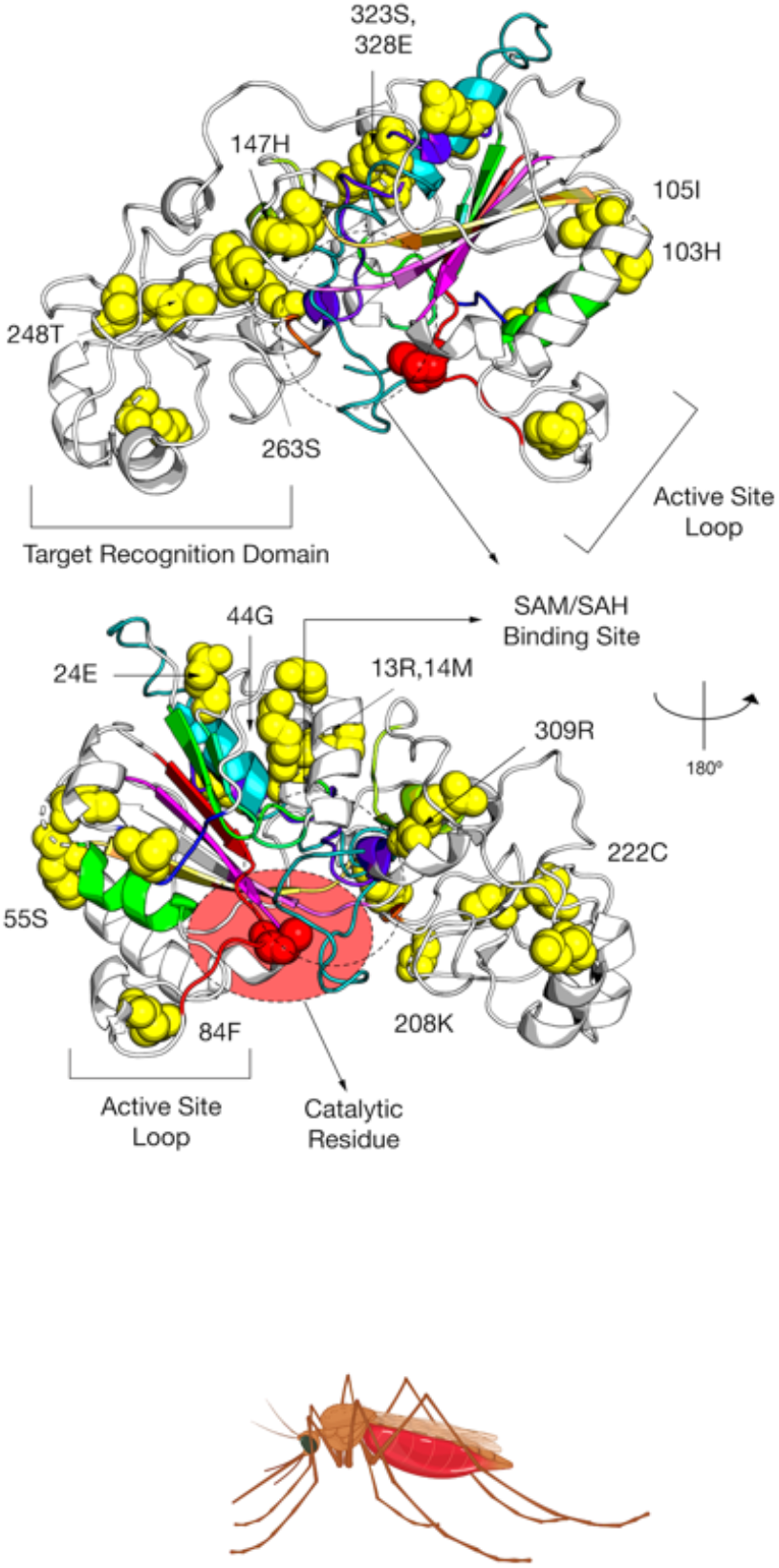
Amino acid sites under positive selection in *Anopheles*. Amino acid positions with evidence of positive selection and high posterior probability values (> 95%) were identified within DNMT2 orthologs from mosquitoes (Culicidae, Figure 1A, Table 1). Spatial distribution of 16 sites unique to *Anopheles darlingi*, which include sites present in ancestral branches (3,19,20,21, Table 1) are represented as yellow spheres on ribbon model of *Anopheles darlingi* DNMT2 structure visualized in PyMOL 2.4 (Schrödinger, LLC). Catalytically active cysteine residue (Cys, C) is represented in red. Predicted substrate i.e. S-adenosyl methionine (SAM) or S-adenosyl homocysteine (SAH) binding sites are indicated in oval with dashed outline. Functionally important active-site loop and target recognition domain are also indicated on each structure. The rotation symbol reflects structural features viewed 180° apart along the vertical axis.

**Supplementary Figure 3:**
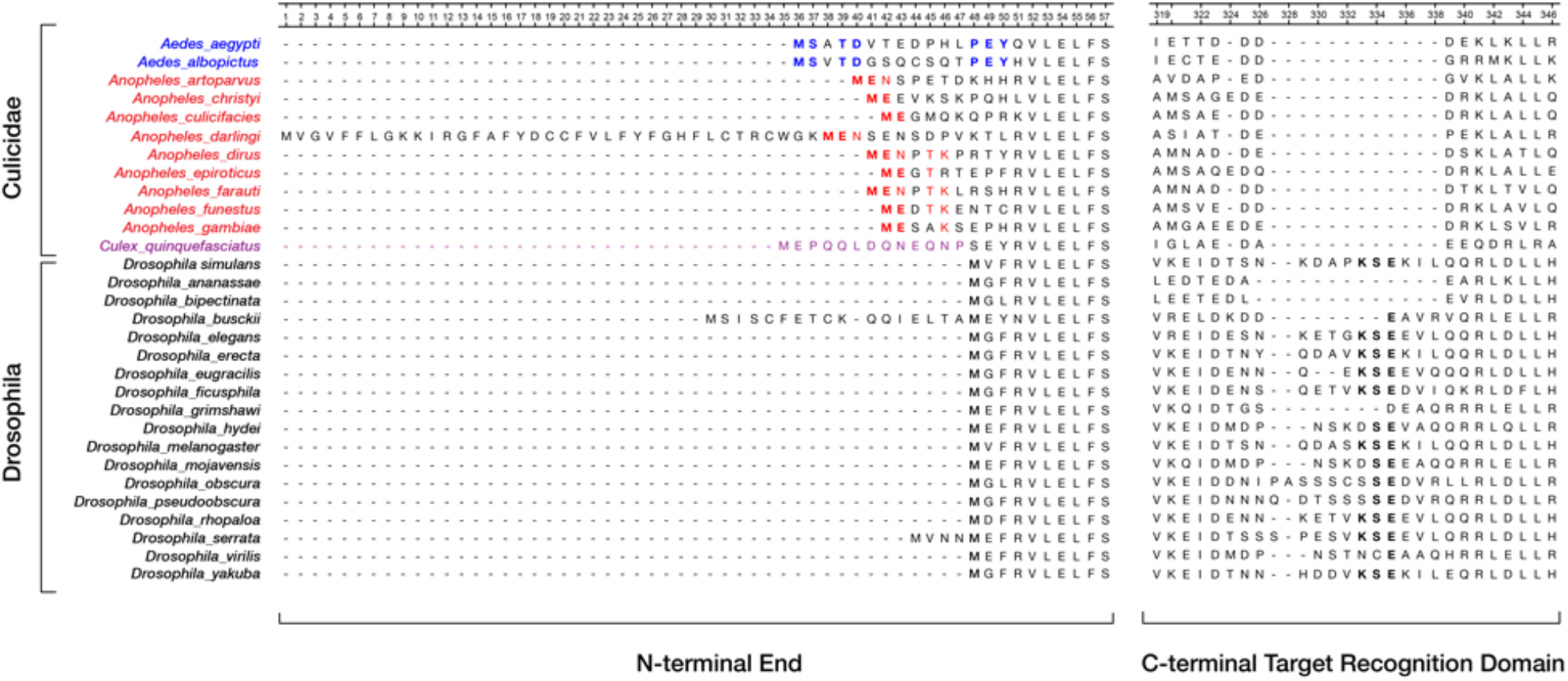
Differences in primary amino acid sequence between Culicidae and Drosophila DNMT2 orthologs. Conserved amino acids present in the extended N-terminal end and the C-terminal Target Recognition Domains of Culicidae Drosophila species represented in a multiple sequence alignment, with Aedes genera in blue, Anopheles genera in red and Culex in purple. Fully conserved residues are in represented in bold letters.

**Supplementary Figure 4:**
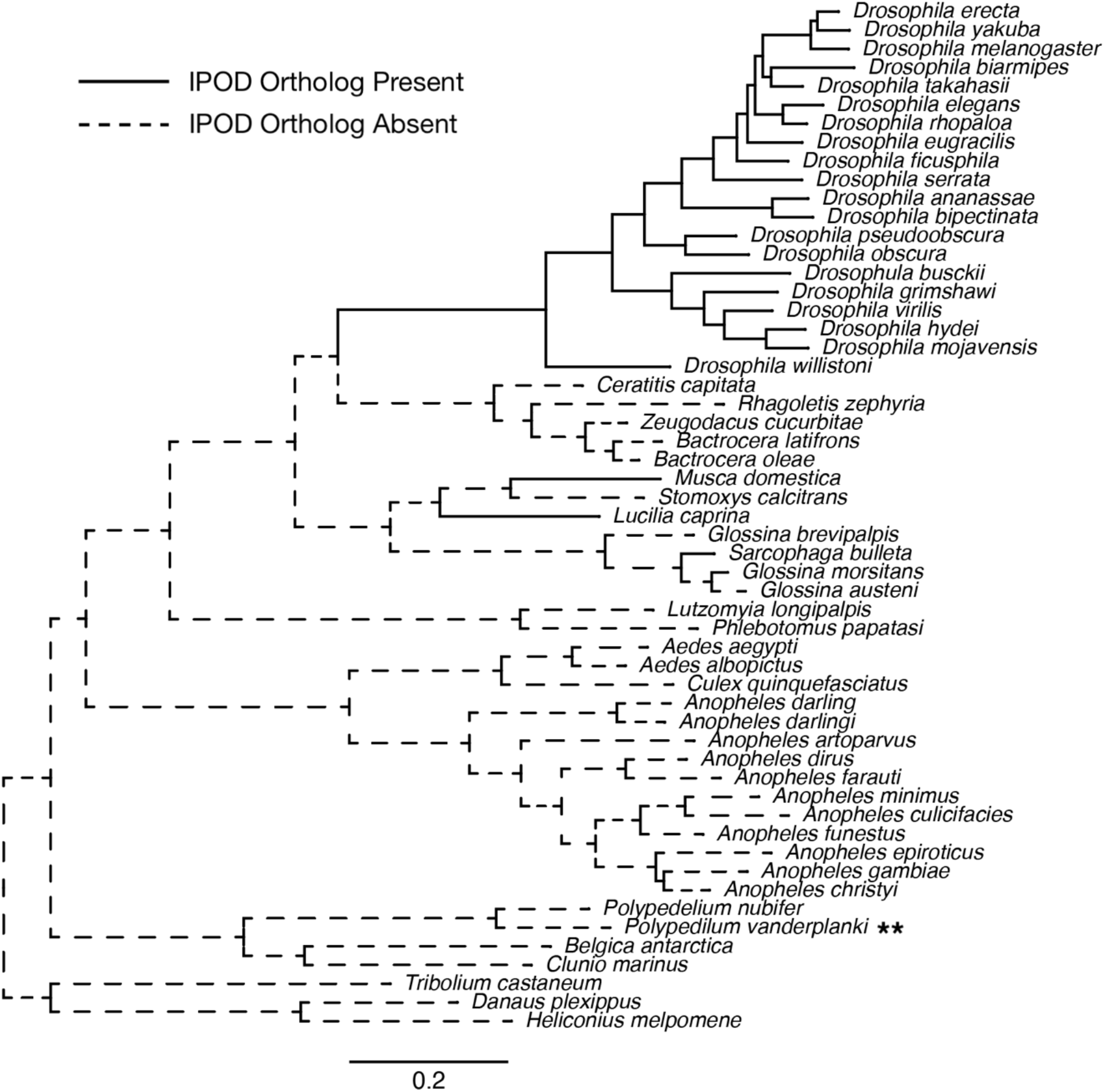
Presence of IPOD orthologs across *Dipterans*. *Dipteran* species with known IPOD orthologs (Protein-BLAST) represented on Maximum-likelihood (Deddouche, et al.) tree constructed using DNMT2 sequences in RAxML. As presented in Figure 4B, IPOD orthologs are present within *Drosophilidae* and only three other *Dipteran* species (represented in this tree in solid branches). Protein-BLAST failed to identify any potential IPOD orthologs in other *Dipteran* species (represented with dashed branches). Taxa label with accompanying asterisks (**) represent the lack of a full genome assembly and therefore should not be considered while interpreting the results. Scale bar represent branch lengths.

**Supplementary Figure 5:**
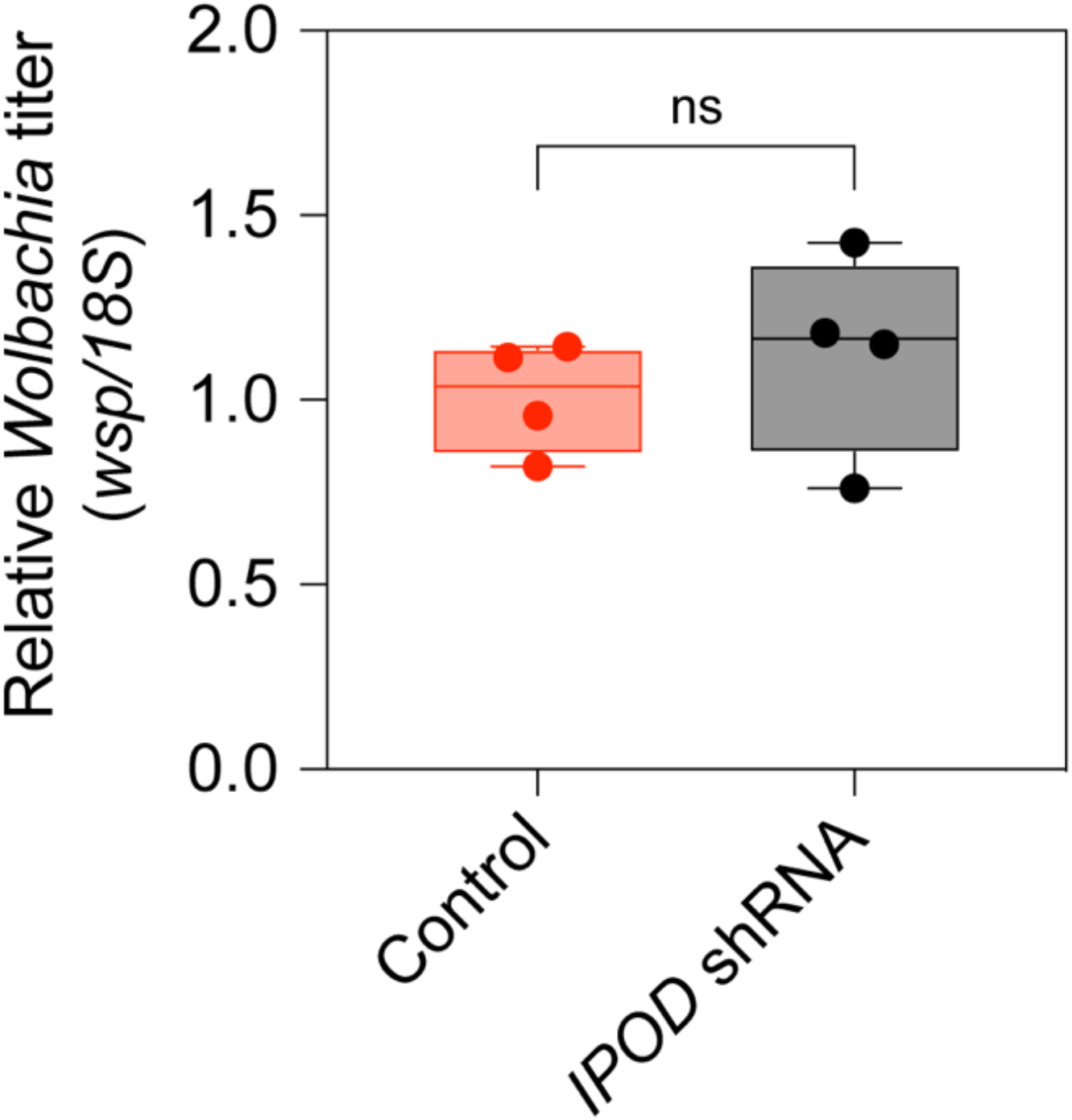
Relative *Wolbachia* titer in *IPOD* knockdown flies. Quantitative RT-PCR was used to measure expression of the *Wolbachia wsp* gene relative to the endogenous ribosomal 18S RNA in age-matched 2-4 days old adult female flies following RNAi-mediated knockdown of *IPOD. IPOD* expression was knocked down in *Wolbachia w*Mel-colonized *Drosophila melanogaster* (TRiP line# 60092) by driving expression of a targeting short-hairpin RNA (shRNA) against the target *IPOD* mRNA. Controls represent isogenic sibling flies not expressing the targeting shRNA. Unpaired Welch’s t-test: p = 0.4788, t = 0.7695, df = 4. Error bars represent standard error of mean of 4 independent experimental replicates. ns = not-significant.

**Supplementary Figure 6:**
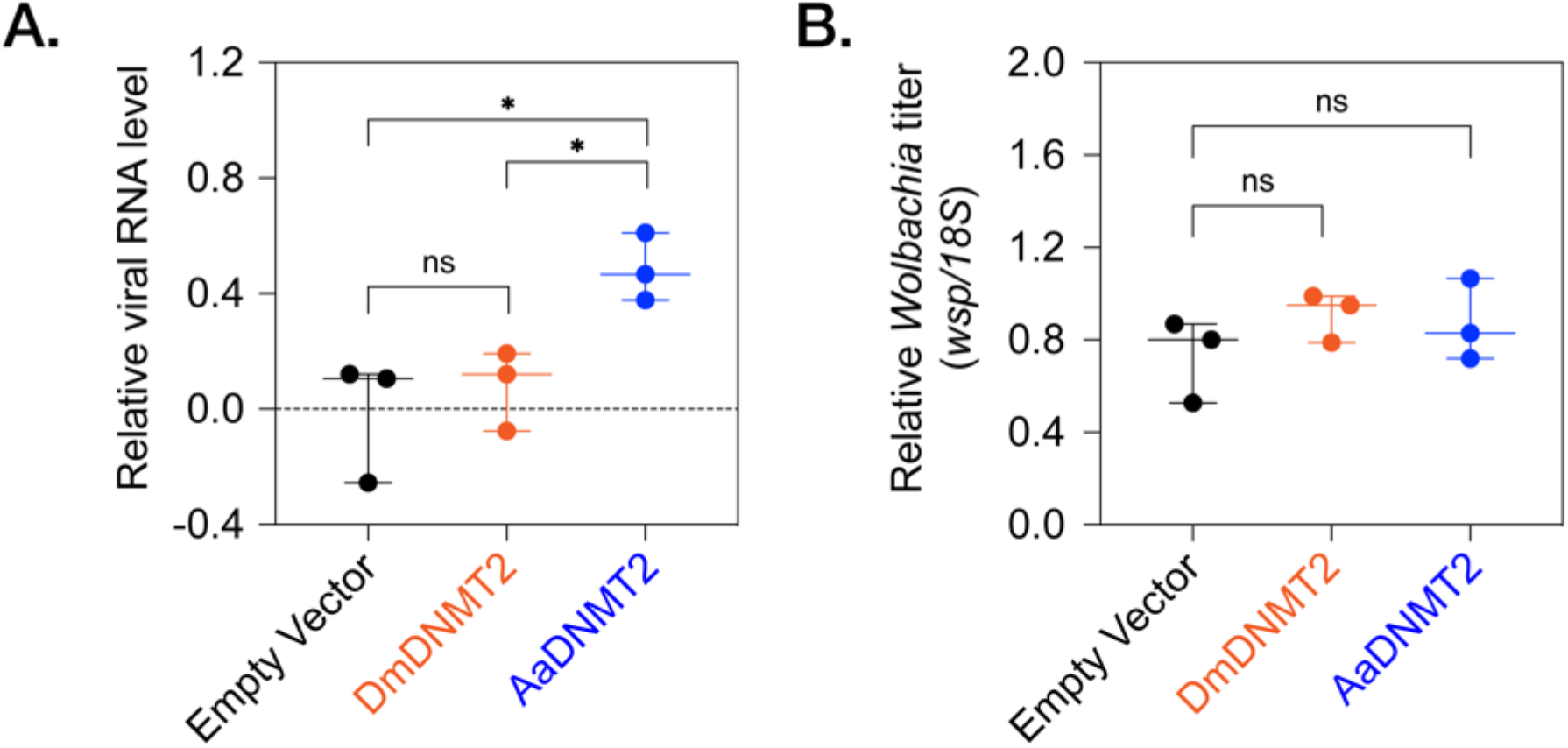
Effect of heterologous DNMT2 expression on *Wolbachia* in mosquito cells. 72 hours post transfection, *Aedes albopictus* derived C710 cells (colonized with *w*Stri Wolbachia strain) expressing either the empty vector, the native DNMT2 (*Dm*DNMT2) or the non-native DNMT2 (*Aa*DNMT2) were challenged with SINV at MOI of 10 particles/cell. Cell lysates were collected 48 hours post infection and levels of (A) virus and (B) *Wolbachia* RNA levels were assessed using qRT-PCR on total extracted RNA. One-way ANOVA with Tukey’s post hoc test for multiple comparisons: SINV RNA, Empty Vector vs *Dm*DNMT2: p = 0.7875, Empty Vector vs *Aa*DNMT2: p < 0.05, *Dm*DNMT2 vs *Aa*DNMT2: p < 0.05, *Wolbachia*, Empty Vector vs *Dm*DNMT2: p = 0.4121, Empty Vector vs *Aa*DNMT2: p = 0.5639, *Dm*DNMT2 vs *Aa*DNMT2: p = 0.9523. Error bars represent standard error of mean of 3 independent experiments. *p < 0.05, ns = not-significant.

**Supplementary Figure 7:**
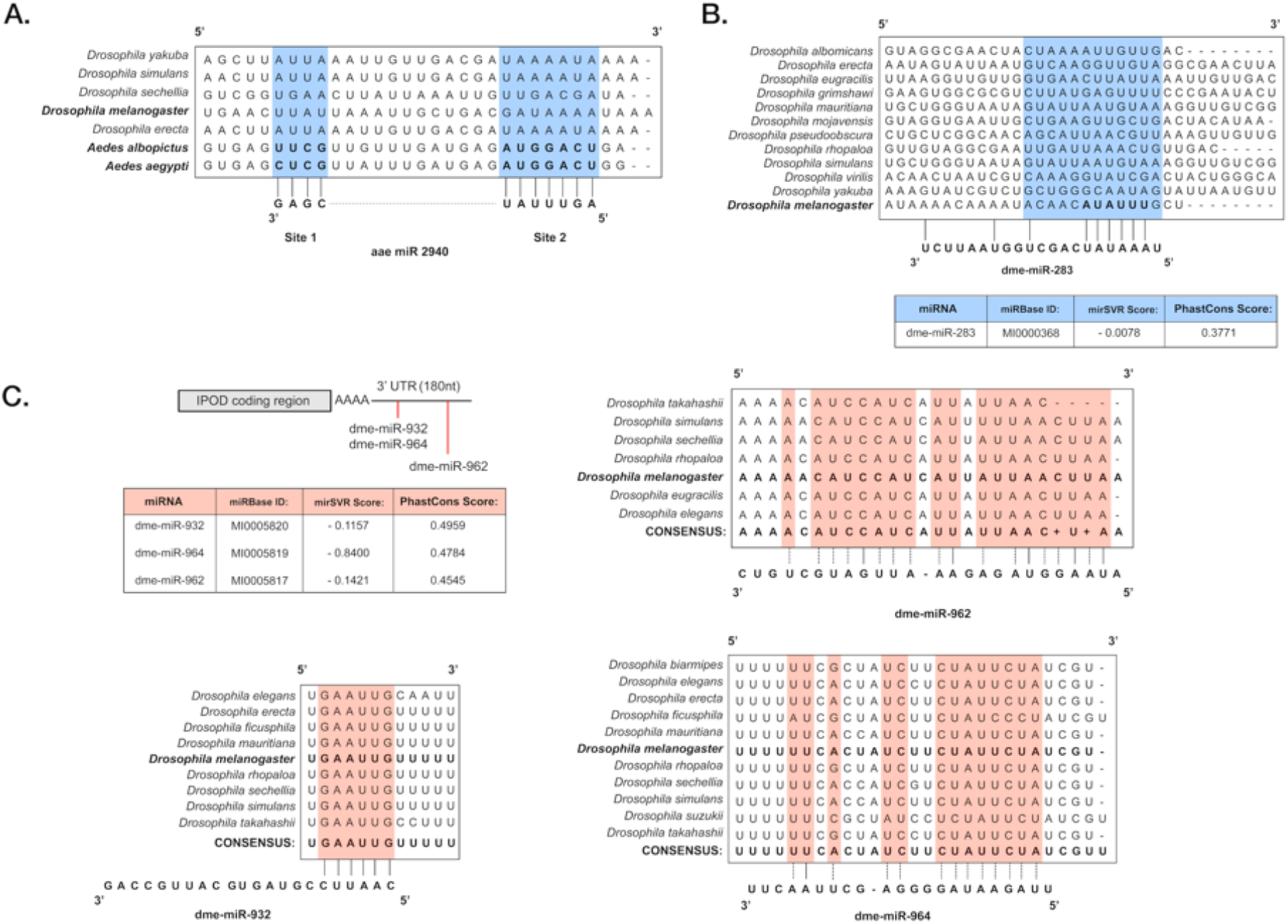
Predicted role of miRNAs in regulation of *Drosophila* DNMT2 and IPOD. (A) Prior studies demonstrate the role of the *Aedes* miRNA *aae-miR-2940-5p* in regulating the expression of *Aedes* DNMT2 orthologs (Zhang et.al. 2013). Target sequence for this miRNA is conserved at the primary nucleotide sequence level in *Aedes albopictus* and *Aedes aegypti* DNMT2 *(Mt2)* mRNAs (indicated in the multiple sequence alignment), but absent in *Mt2* mRNAs of orthologs present in all *Drosophila* species, including *Drosophila melanogaster* (taxa depicted in bold letters). (B) Location of conserved miRNA dme-miR-283 predicted to target the 3’ untranslated region (3’UTR) of *Drosophila melanogaster* DNMT2 (taxon depicted in bold letters). (C) Location of conserved miRNAs predicted using mirSVR (microrna.org) to target the 3’ untranslated region (3’UTR) of IPOD. Empirical probability of target downregulation for each miRNA, considering the conservation of the target site, is indicated by the mirSVR downregulation scores. Evolutionary conservation of each of the miRNA target sequences is indicated by the PhastCons scores (PHylogenetic Analysis with Space/Time models CONServation). (C) Sequence conservation of miRNA target region(s) are depicted in light red within aligned nucleotide sequences of 3’UTR regions belonging to IPOD orthologs of different *Drosophila* species. Nucleotide sequence(s) of *Drosophila melanogaster* IPOD is depicted in bold letters.

**Supplementary Table 1.**
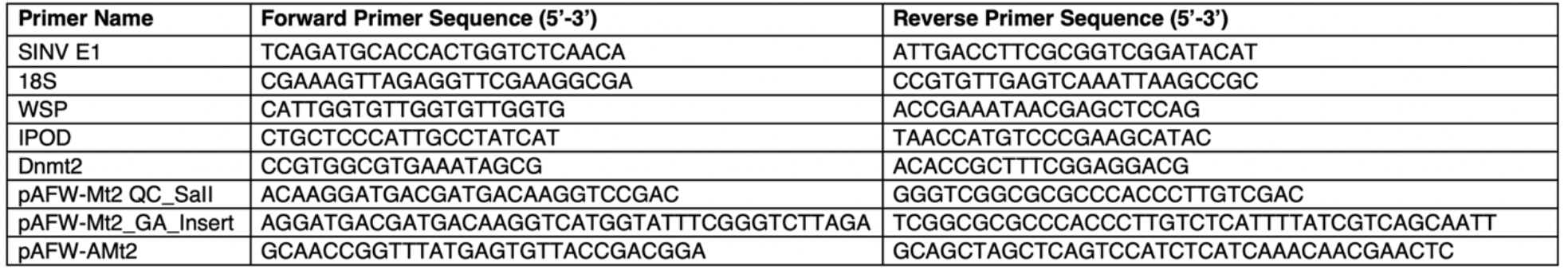
Primers used in this study. Primers were purchased from Integrated DNA Technologies (IDT). All primers were used at a final concentration of 10μM for PCR and quantitative RT-PCR reactions.

